# Inferring Protein Variant Impacts Across Contexts

**DOI:** 10.64898/2026.08.18.745369

**Authors:** Alireza Rasoulzadeh, Vignesh Senguttuvan, Warren van Loggerenberg, Richard Border, Frederick P. Roth

**Affiliations:** Department of Computational and Systems Biology, School of Medicine, University of Pittsburgh, 3396 Fifth Avenue, 10th floor, Pittsburgh, PA 15213, United States; Ray and Stephanie Lane Computational Biology Department, School of Computer Science, Carnegie Mellon University, Pittsburgh, PA 15213, United States

## Abstract

Multiplexed assays of variant effects (MAVEs) measure the functional impact of many protein sequence variants in parallel, potentially covering all possible single amino acid substitutions. Unlike current computational variant effect predictors, MAVEs can reveal the effects of variants under different genetic and environmental contexts. However, whereas the space of possible contexts is effectively infinite, ‘contextual’ MAVE studies are limited by finite experimental budgets. To maximize coverage across contexts, one strategy is to carry out sub-saturation contextual MAVEs and then fill in the gaps via imputation. Here we categorize and compare different imputation challenges, explore a collection of multi-context imputation solutions (including linear mixed-effects models, random forests, and autoencoders), and provide insight into how best to proceed in any given imputation task. We find that which method is optimal depends on the imputation task and on how densely the contexts have been measured, with more flexible models excelling when measurements are plentiful and the simplest models proving most reliable when measurements are sparse. However, the simplest models, though well suited to imputing scores where variants have been measured in the source context, cannot impute scores where variants were not measured in either context. This proves to be a major limitation when both maps are sparsely measured. We provide a conceptual framework and an initial evaluation of multi-context imputation methods that can extend the scope of large-scale studies of context-dependent variant effects.

## Introduction

Despite the rapidly increasing catalog of human genetic variation and its importance to disease (Shendure and Akey 2015), our ability to identify which rare sequence variants have a disease-causing functional impact remains limited. Fewer than 2% of observed missense variants have been clinically classified as pathogenic or benign (Cheng et al. 2023). Moreover, variants of uncertain significance (VUSs) constitute the largest single classification category in ClinVar (Fowler and Rehm 2024; Starita et al. 2017). This ‘VUS crisis’ hampers diagnosis and therapy while burdening patients with anxiety. The scale of the pathogenicity classification challenge is underscored by a population-scale estimate that, if genome sequences were available for everyone alive today, each single-nucleotide variant compatible with life would appear about 50 times on average (Wu et al. 2021). A further complication is that variant effects are context-dependent. Ultimately, clinicians and their patients want to know whether a variant has caused or will cause disease in a specific person given their individual genetic background and environmental exposures.

Evidence of variant functional impact to inform clinical classification can come from two key sources. First, variant effect predictors (VEPs) are in silico models that can predict pathogenicity (Grimm et al. 2015; Livesey and Marsh 2025; Tabet et al. 2024). A valuable aspect of VEPs is that most of the current top predictors offer pre-calculated scores for nearly all possible missense variants for nearly all genes (Cheng et al. 2023; Frazer et al. 2021). Although VEPs have historically been interpreted as offering only the weakest level of evidence towards variant classification (Richards et al. 2015), more recent calibrations have shown that some tools can provide moderate or strong evidence at calibrated score thresholds (Pejaver et al. 2022; Bergquist et al. 2025). However, current VEPs provide a single static score for each variant, without awareness of dependence of variant impacts on environmental and genetic contexts, and often without providing insights into underlying mechanisms (Livesey and Marsh 2025; Grimm et al. 2015; Wu et al. 2021).

The second key source of evidence for variant functional impacts is multiplexed assays of variant effects (MAVEs), a powerful and scalable set of experimental technologies that can uncover sequence-function relationships and classify clinical variants (Starita et al. 2017; Tabet et al. 2022). Many MAVEs have been shown to perform well in distinguishing pathogenic from benign variants (Findlay et al. 2018; Weile et al. 2021; Gebbia et al. 2024; Livesey and Marsh 2025). However, MAVEs have important limitations: they are typically performed one protein at a time, often require bespoke experimental designs, and are resource-intensive. Existing variant effect maps cover less than 1% of the known clinically relevant human genome (Fowler et al. 2023). However, and of particular importance for this study, MAVEs can not only reveal variant functional impacts, but can also systematically reveal how those impacts depend on environmental and genetic context.

Even where MAVEs are carried out in a single context, it can be challenging to achieve saturation with confident measurements provided for all possible nucleotide or amino acid substitutions. For this reason, several groups have developed within-map (W) imputation strategies to fill in missing variant scores (or refine less confidently measured values) without requiring a separate source-context map (Weile et al. 2017; Yu et al. 2024; Polunina et al. 2026; Wu et al. 2019; Zhou et al. 2022). Although imputation can incorporate patterns of conservation or other features used by current-generation variant effect predictors to generate more accurate impact scores, evidence for variant classification based purely on experimental measurements may be more reasonably treated as independent from other types of variant effect predictors (Gebbia et al. 2024).

Although measuring variant effects using multiple assay types or under different environmental or genetic settings can provide additional insights into mechanism and context-dependence of variant effects, it is impracticable to deploy saturated MAVEs for all disease-associated proteins under all relevant assay, genetic and environmental contexts. However, the scope of MAVEs can be extended by inferring fully saturated variant effect maps from a collection of sub-saturation contextual variant effect maps, via cross-context imputation.

We explored two classes of imputation. The first is the W imputation approach described above. The second, more novel, class is between-map (B) imputation, which uses a separate source map to predict scores in a target map, and further divides by whether the variant of interest was measured in the source: the source-informed case (B₁), where the source score is available, and the missing-source case (B_0_), where the variant is missing in both source and target. Both W and B methods were evaluated by their ability to reconstruct held-out variant effects within a MAVE dataset collected under multiple genetic and environmental contexts. Together, our results provide methods and guidance for cross-context imputation. These methods collectively enable, given a fixed experimental budget, a deeper exploration of genetic and environmental contexts while still yielding highly saturated knowledge of variant functional impacts.

## Material and methods

### MTHFR maps and their use to evaluate imputation methods

Scores from eight 5,10-methylenetetrahydrofolate reductase (MTHFR) variant effect maps (Weile et al. 2021), encompassing four different folinate concentration contexts in both the presence and absence of the p.A222V context, were derived from variant counts files generated by aligning paired-end reads to a reference sequence using Bowtie2 (Langmead and Salzberg 2012). Variants were called based on Phred scores achieving a posterior probability of 90% or greater. Variant counts were tabulated, normalized according to sequencing depth, and used to calculate marginal variant frequencies. Quality filtering retained scores for variants with sufficient counts in the ‘non-select’ condition (not subjected to selection for variant function) and removed scores where disagreement between replicates exceeded that observed for wild-type (WT) controls. Enrichment values were calculated as log ratios of select to non-select variant frequencies, and experimental error was estimated as described (Weile et al. 2017). Finally, fitness scores were linearly rescaled such that the medians of the nonsense and synonymous distributions were 0 and 1, respectively.

We assessed imputation performance using Monte Carlo cross-validation and evaluated root mean square error (RMSE) on held-out entries as our primary accuracy metric. We define saturation as the fraction of observed scores retained (uniformly at random) for training (for example, 60% saturation corresponds to holding out 40% of observed scores as test data). Performance was evaluated at six saturation levels: 90%, 80%, 60%, 40%, 20%, and 10%. For each saturation level, 50 random splits were generated by masking observed scores completely at random (MCAR) (Rubin 1976), and RMSE was computed on the held-out values.

We evaluated two imputation strategies: within-map (W) imputation, which predicts missing scores without using measurements from a separate source-context map, and between-map (B) imputation, which uses a source map to predict scores in a target map. Here, a map denotes a single MAVE measurement context (e.g., one folinate concentration in one genetic background). For between-map evaluations, we define a source map (the context from which we extrapolate) and a target map (the context where we predict missing scores and compute RMSE).

For between-map imputation (B), we distinguish two sub-tasks based on the availability of source information. In the source-informed case (B₁), the variant’s score was measured in the source map but was missing in the target map, allowing the imputer to leverage the source observation directly. In the missing-source case (B_1_), the variant’s score is missing in both maps. Because B_0_ is substantially harder than B₁, and because several models cannot address it, results for the two sub-tasks were reported separately.

### Baseline method comparison

For both within and between-map imputation, all methods were compared against a Column Mean baseline method: for each amino acid residue position, every missing score was imputed as the arithmetic mean of all measured scores at that position. Where no measured scores were available at a given position, imputation was based on the global mean across all positions.

### Within-map imputation approaches

In addition to the baseline approach, we applied four additional method types for within-map (W) imputation. The first was a k-nearest neighbors imputer in which distance was based on amino acid substitutability (kNN-BLOSUM) with k = 5. For each missing score, the BLOSUM100 substitution score (Henikoff and Henikoff 1992) between the mutant amino acid and each candidate amino acid at the same position was computed, and the k = 5 candidates with the highest scores were selected as nearest neighbors. The imputed value was taken as the mean of the subset of scores available for these k nearest substitutions. Where 0 or 1 of these scores were available at a given position, the position mean (or global mean if no position scores were available) was used as a fallback. Nonsense (Ter) mutations were handled separately using scores from nonsense mutations at neighboring positions.

The second was principal-component analysis (PCA)-based imputation. This was based on the set of vectors (one for each of the 656 amino acid positions in MTHFR), each vector having 22 scores (one for each of 20 possible substitutions, plus synonymous and nonsense). An iterative expectation-maximization procedure was used. Missing values were initialized with per-substitution column means, PCA was fit to the completed matrix, missing values were updated with PCA-reconstructed values, and the process was repeated until convergence (maximum change in imputed values below 1 × 10⁻⁴, up to 20 iterations). We tested k = 1, 2, 4, 10, 20, and 22 principal components; k = 1 was used as the representative configuration in the main results, with sensitivity analysis across k values presented in Supplemental Figure S1.

The third was Multiple Imputation by Chained Equations with a random forest estimator (MICE-RF). The model was provided with a table of scores from a single context along with covariates including amino acid position, wild-type amino acid identity, mutant amino acid identity, and protein domain annotation. Each residue position with missing values was treated as a response variable to be predicted from the scores at all other positions and the available covariates, with the procedure iterated until convergence. The resulting completed dataset was used to compute RMSE on held-out entries (Wilson 2021; van Buuren and Groothuis-Oudshoorn 2011).

The fourth was a masked autoencoder, SingleAE (within-map): a fully connected encoder-decoder network trained to reconstruct missing entries in the score matrix. The autoencoder was provided with a set of 22-dimensional vectors as described above for PCA. The encoder mapped partially observed substitution profiles into a low-dimensional latent representation that captured shared structure across positions, while the decoder reconstructed the full 22-dimensional vector. Training loss was computed exclusively over observed entries (masked mean squared error [MSE]), so the network was not penalized for predictions at unobserved positions (Vincent et al. 2008). After training, decoder outputs at originally missing indices provided the imputed fitness scores.

### Between-map imputation methods

We applied five method types for between-map (B) imputation, ordered below from simplest to most complex.

For the first method type, we evaluated both fixed and mixed-effects regression approaches. The fixed-effects models incorporated features including source map score, protein domain annotations, and relevant interaction terms. The mixed-effects models extended this framework by introducing random effects to account for structured variation across amino acid identities. Both formulations are applicable to the B₁ scenario only, as they require an observed source score as input. Five model specifications were evaluated: Basic Linear, 1-Param Nonlinear, a domain-specific model allowing slope and intercept to vary by protein domain (Linear + Domain), a mixed random-intercept model with domain interaction in the fixed effects and amino acid identity as a random intercept (Mixed (rand. int.)), and a mixed model with random slope and intercept by amino acid identity (Mixed (rand. slope)).

The second was MICE with predictive mean matching (MICE-PMM). Each imputation dataset consisted of scores from the source and target maps along with covariates including amino acid position, protein domain, wild-type amino acid, and mutant amino acid identity. MICE-PMM fits a regression model at each step, then imputes missing values by selecting from observed donor values whose predicted values are closest to the predicted value for the missing entry, rather than using the regression prediction directly (van Buuren and Groothuis-Oudshoorn 2011). This approach ensures that imputed values are always drawn from the observed data distribution. This method addresses both the B_1_ and B_0_ scenarios, as the chained equations procedure can generate predictions for entries missing in both source and target maps.

The third was MICE with a random forest estimator (MICE-RF), as implemented using the miceRanger package (Wilson 2021). This method follows the same chained imputation framework as MICE-PMM but substitutes a random forest model at each step, thereby capturing nonlinear dependencies among maps that a linear estimator cannot. Unlike MICE-PMM, the random forest variant uses the model’s predicted value directly rather than selecting from a donor pool. Like MICE-PMM, this method addresses both the B_1_ and B_0_ scenarios.

The fourth was a masked single autoencoder for between-map prediction (SingleAE (cross-map)). Here we used the same encoder-decoder architecture as for SingleAE within-map imputation, but instead trained it to predict target map scores from source map scores. For each amino acid position, the 22-dimensional source vector was passed through the network to produce a 22-dimensional target prediction. The network was trained using masked MSE loss computed over positions where both source and target scores were observed. At inference, predictions were inserted at positions missing in the target map, with observed target values retained. Since the encoder processes the full source vector at each position regardless of individual component availability, this method addresses both the B_1_ and B_0_ scenarios.

The fifth was a dual-decoder masked autoencoder (DualAE). This was an extension of SingleAE (cross-map) with two output branches: one reconstructing the source map and one predicting the target map. By simultaneously reconstructing source and predicting target scores, the encoder learns a latent representation that captures cross-context structure even in the absence of direct source observations. The combined training loss was a weighted sum of masked reconstruction loss on the source map (weight 0.7) and masked prediction loss on the target map (weight 0.3). Because the model does not require the source score at a given position to generate a target prediction, it addresses both the B_1_ and B_0_ scenarios.

## Results

To evaluate the performance of different imputation strategies, we benchmarked the methods using data for the MTHFR protein. MTHFR encodes a key enzyme in one-carbon metabolism and folate processing (Blomgren et al. 2024). Severe MTHFR deficiency causes hyperhomocysteinemia and can affect the central nervous system and produce thromboembolic, ocular, and other multisystem manifestations (Huemer and Baumgartner 2019; Rommer et al. 2017). Because some patients can be treated with dietary supplementation for folate deficiency, a study by Weile et al. (2021) systematically measured variants in the context of each of four different folinate concentrations (folinate and folate are rapidly interconverted with each other and with MTHFR’s substrate). Moreover, because the common MTHFR p.A222V variant reduces enzymatic activity and its fitness defect worsens as folinate availability decreases (Liew and Das Gupta 2015; Weile et al. 2021), Weile et al. also studied variant effects across the same four folinate concentration contexts in the presence of the p.A222V variant. This yielded eight variant effect maps spanning four folinate concentrations in two genetic backgrounds (wild-type and p.A222V), providing an ideal testbed to evaluate imputation across contexts that differ in both environmental and genetic dimensions.

### Imputation tasks overview

We explored two classes of imputation problems: within-map (W) imputation, which predicts missing scores without using measurements from a separate source-context map, and between-map (B) imputation, which uses a source map to predict scores in a target map. The latter can be further divided into missing-source (B_0_) and source-informed (B_1_) tasks. Figure 1 summarizes these task definitions and the autoencoder architectures.

**Figure 1.**
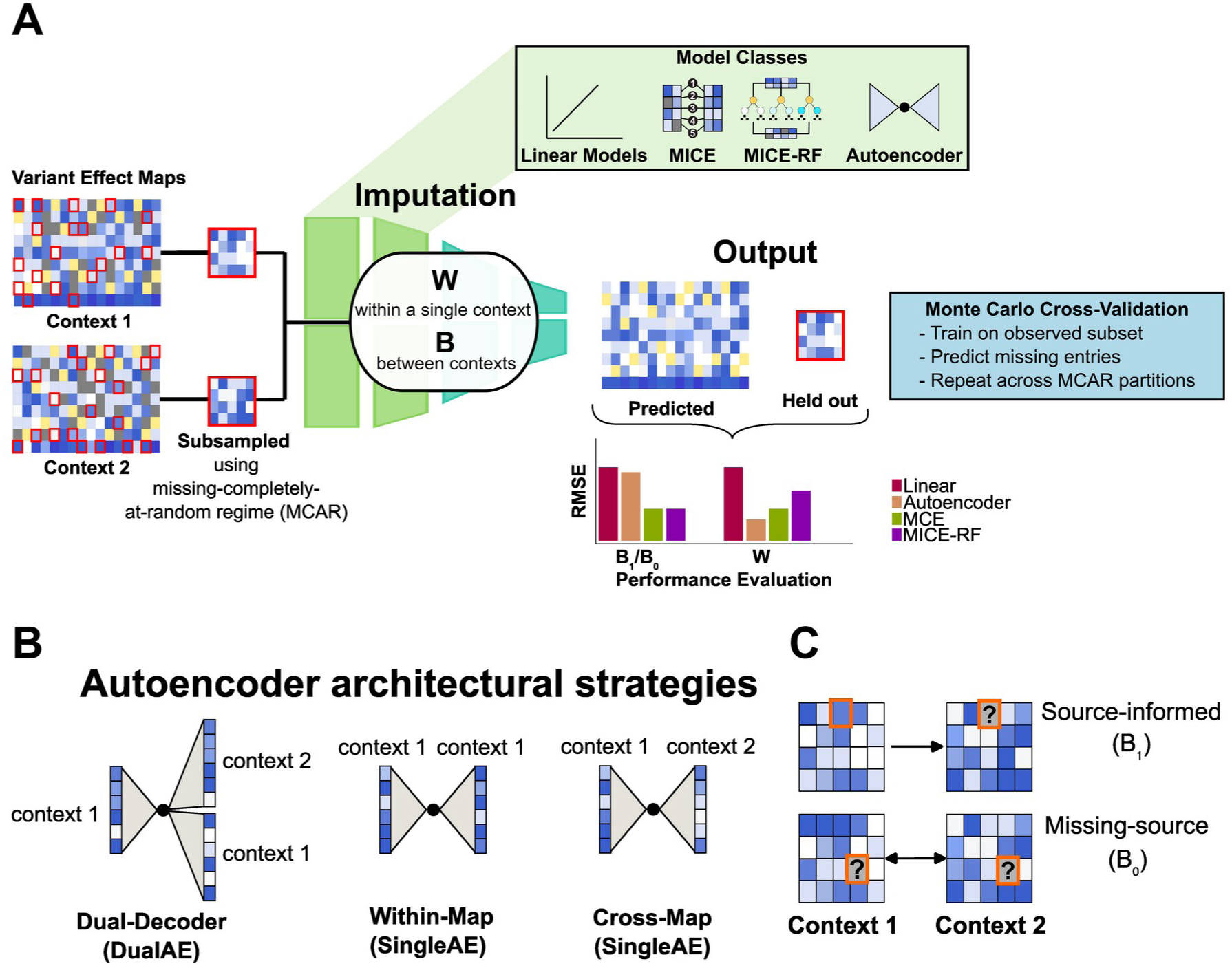
Within-map and between-map imputation workflow. (A) Workflow used to benchmark imputation of variant effect maps. Originally observed scores were randomly held out in each of 50 Monte Carlo splits. Predictions from different models were compared with held-out target values using RMSE. Within-map (W) imputation uses measurements from one context, whereas between-map (B) imputation uses measurements across contexts. (B) Autoencoder architectures. SingleAE (within-map) reconstructs the input map, SingleAE (cross-map) predicts a target map from a source map, and DualAE uses a shared encoder with separate source-reconstruction and target-prediction decoders. (C) Between-map held-out values are classified as source-informed (B_1_) when the corresponding source score is available and missing-source (B_0_) when it is unavailable. RMSE is calculated only for held-out target values that were measured in the original data.

We evaluated all three scenarios (W, B_1_, B_0_) using Monte Carlo cross-validation. For each of six saturation levels (10%, 20%, 40%, 60%, 80%, 90%) we generated 50 random splits that held out a fraction of observed scores and computed RMSE on the held-out values. Performance was compared against a Column Mean baseline across all applicable models (see Methods). Predictive accuracy on held-out target values is summarized in Table 1 (between-map) and Table 2 (within-map).

**Table 1.**
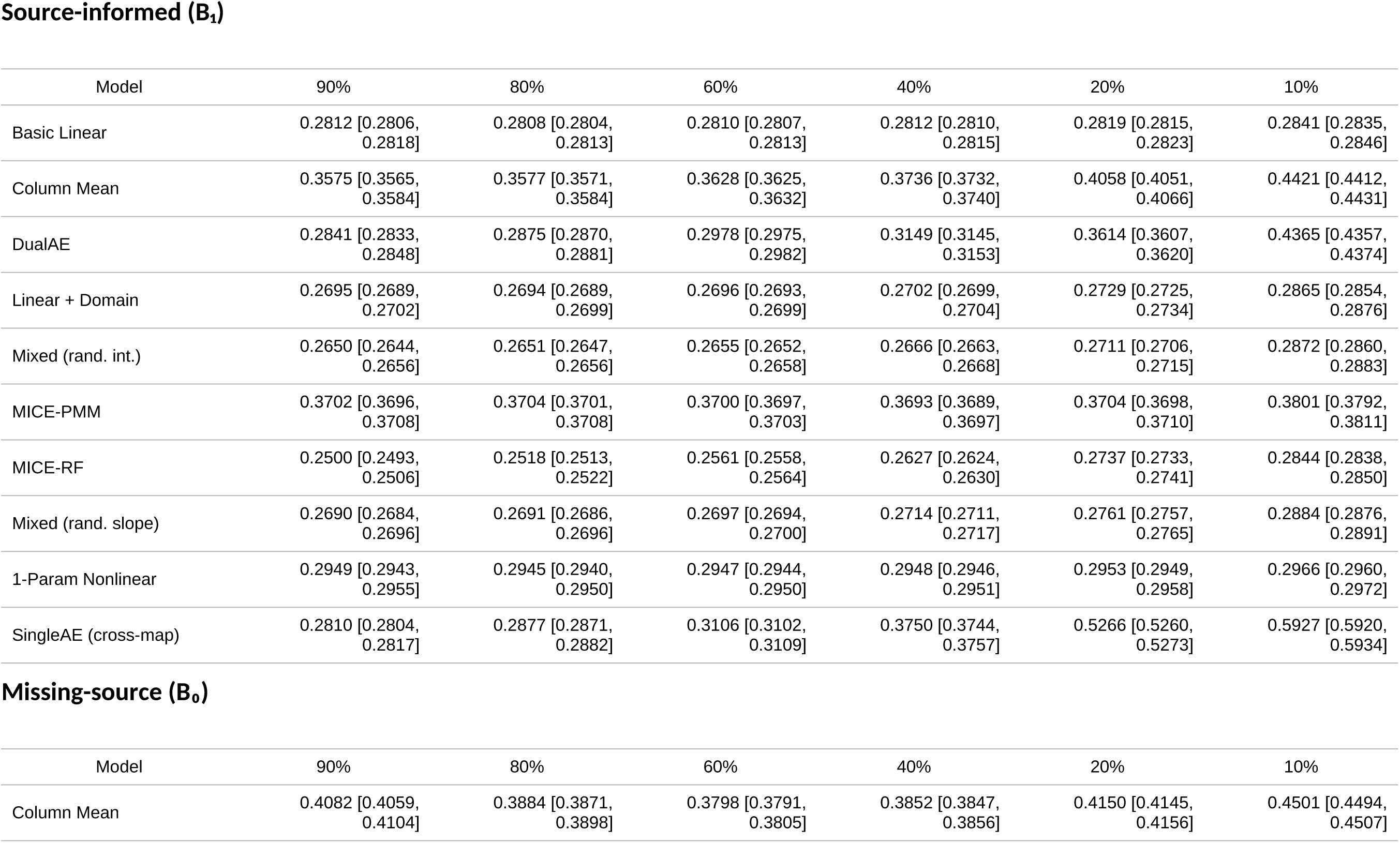
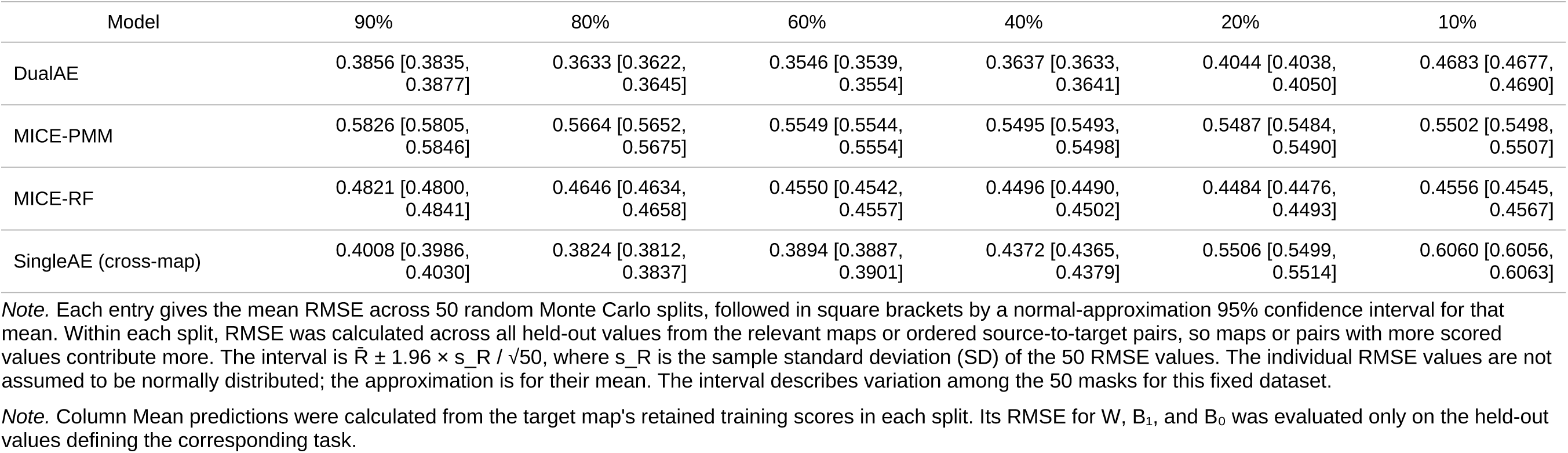
RMSE on held-out values for source-informed and missing-source prediction.

**Table 2.** RMSE on held-out values for within-map imputation.

Within-map (W)
| Model | 90% | 80% | 60% | 40% | 20% | 10% |
| --- | --- | --- | --- | --- | --- | --- |
| Column Mean | 0.3649 [0.3639, 0.3659] | 0.3652 [0.3645, 0.3658] | 0.3702 [0.3697, 0.3706] | 0.3808 [0.3805, 0.3811] | 0.4133 [0.4127, 0.4138] | 0.4493 [0.4487, 0.4500] |
| kNN-BLOSUM | 0.3389 [0.3379, 0.3398] | 0.3431 [0.3425, 0.3438] | 0.3517 [0.3513, 0.3521] | 0.3634 [0.3631, 0.3637] | 0.3949 [0.3944, 0.3955] | 0.4312 [0.4306, 0.4319] |
| MICE-RF | 0.3669 [0.3660, 0.3678] | 0.3672 [0.3666, 0.3678] | 0.3690 [0.3686, 0.3694] | 0.3732 [0.3729, 0.3736] | 0.3803 [0.3800, 0.3807] | 0.3893 [0.3889, 0.3898] |
| PCA (k = 1) | 0.3271 [0.3262, 0.3280] | 0.3280 [0.3273, 0.3287] | 0.3340 [0.3335, 0.3345] | 0.3473 [0.3470, 0.3476] | 0.3857 [0.3851, 0.3862] | 0.4606 [0.4586, 0.4626] |
| SingleAE (within-map) | 0.3651 [0.3641, 0.3661] | 0.3649 [0.3642, 0.3655] | 0.3703 [0.3698, 0.3708] | 0.3826 [0.3822, 0.3830] | 0.4113 [0.4108, 0.4118] | 0.4633 [0.4627, 0.4639] |

Note. Each entry gives the mean RMSE across 50 random Monte Carlo splits, followed in square brackets by a normal-approximation 95% confidence interval for that mean. Within each split, RMSE was calculated across all held-out values from the relevant maps, so maps with more scored values contribute more. The interval is $\bar{R} \pm 1.96 \times s_R / \sqrt{50}$ , where $s_R$ is the sample standard deviation (SD) of the 50 RMSE values. The individual RMSE values are not assumed to be normally distributed; the approximation is for their mean. The interval describes variation among the 50 masks for this fixed dataset.
Note. Column Mean predictions were calculated from the target map's retained training scores in each split. Its RMSE for $W$ , $B_1$ , and $B_0$ was evaluated only on the held-out values defining the corresponding task.

### Within-map imputation: Simple methods provide a strong foundation

For within-map imputation, in which missing scores are predicted without measurements from a separate source-context map, we evaluated five methods: a kNN imputer based on BLOSUM amino acid similarity (kNN-BLOSUM), a compact autoencoder (SingleAE (within-map)), MICE with random forest predictors (MICE-RF), PCA-based reconstruction, and a Column Mean baseline.

At high saturation, all within-map methods were reasonably successful (mean RMSE 0.327– 0.367 at 90% saturation), which shows that this is a broadly tractable imputation task. For context, the root mean square of the experimental error estimates provided by the original MTHFR publication ranged from 0.17 to 0.21 across the 8 MTHFR maps. Among the methods tested, PCA with one component achieved the lowest RMSE from 90% through 40% saturation (0.327 at 90% saturation). kNN-BLOSUM also performed well, likely because it directly exploits known tendencies for one amino acid to functionally substitute for another at a given position (Gray et al. 2017). The Column Mean baseline performed surprisingly well throughout, but did not consistently outperform PCA or SingleAE (within-map). This reflects the fact that position-specific mean scores capture much of the variation in variant impacts within a given map. The complete within-map comparison is shown in Figure 2 and Table 2.

**Figure 2.**
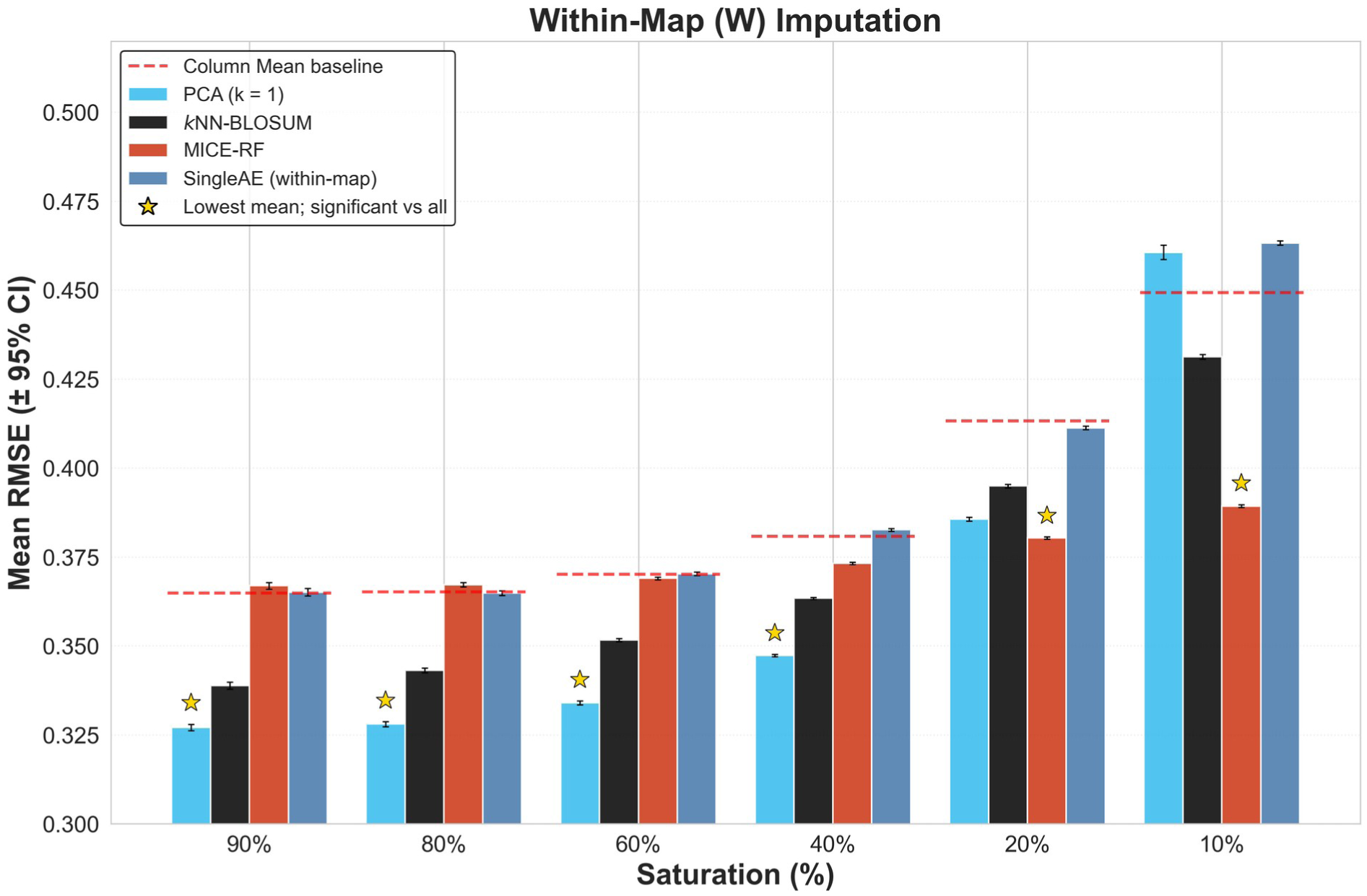
Within-map imputation accuracy across saturation. Within-map RMSE is shown for kNN-BLOSUM, PCA, MICE-RF, and SingleAE (within-map) across six saturation levels. The dashed red line shows Column Mean evaluated on the same W held-out values as the other displayed methods. Bars show mean RMSE on held-out values across 50 random Monte Carlo splits. Error bars show 95% confidence intervals for the mean across splits. A gold star marks the displayed method with the lowest mean only when its two-sided unpaired Mann–Whitney U test against every other displayed method remains significant after Bonferroni correction for that task and saturation. Column Mean was included in these comparisons.

As the fraction of missing data increased, methods diverged in their robustness. MICE-RF degraded the most slowly and, at low saturation, achieved the lowest (best) RMSE (0.389 at 10% saturation), consistent with the random forest’s ability to leverage covariates (amino acid identity, position, protein domain) when training data become scarce. SingleAE was competitive at high saturation but performed worse than the Column Mean baseline at 10% saturation (0.463 vs. 0.449). PCA-based imputation with a single component (rank 1 approximation) performed best from 90% through 40% saturation, but degraded sharply at low saturation and performed worse than the Column Mean at 10% saturation (0.461 vs. 0.449).

These results establish that within-map imputation provided moderate accuracy improvement over a naive baseline approach, but struggled to get below a certain RMSE threshold. The best within-map method achieved an RMSE as low as ∼0.327. As we show below, between-map methods that leverage information from other experimental contexts can achieve substantially lower error when source observations are available (B_1_).

### Between-map source-informed (B_1_) imputation

We next evaluated between-map imputation for the source-informed (B_1_) case. Nine source-informed methods were compared: five linear/mixed models (Basic Linear, 1-Param Nonlinear, Linear + Domain, Mixed (rand. int.), Mixed (rand. slope)), two MICE variants (MICE-PMM and MICE-RF), and two autoencoders (SingleAE (cross-map) and DualAE). The target-only Column Mean reference was evaluated on the same B_1_ held-out values.

MICE-RF achieved the best (lowest) RMSE at high-to-moderate saturation levels (0.250 at 90% saturation, 0.256 at 60% saturation), demonstrating the value of flexible nonlinear modeling when sufficient training data were available. The complete source-informed comparison is shown in Figure 3 and Table 1.

**Figure 3.**
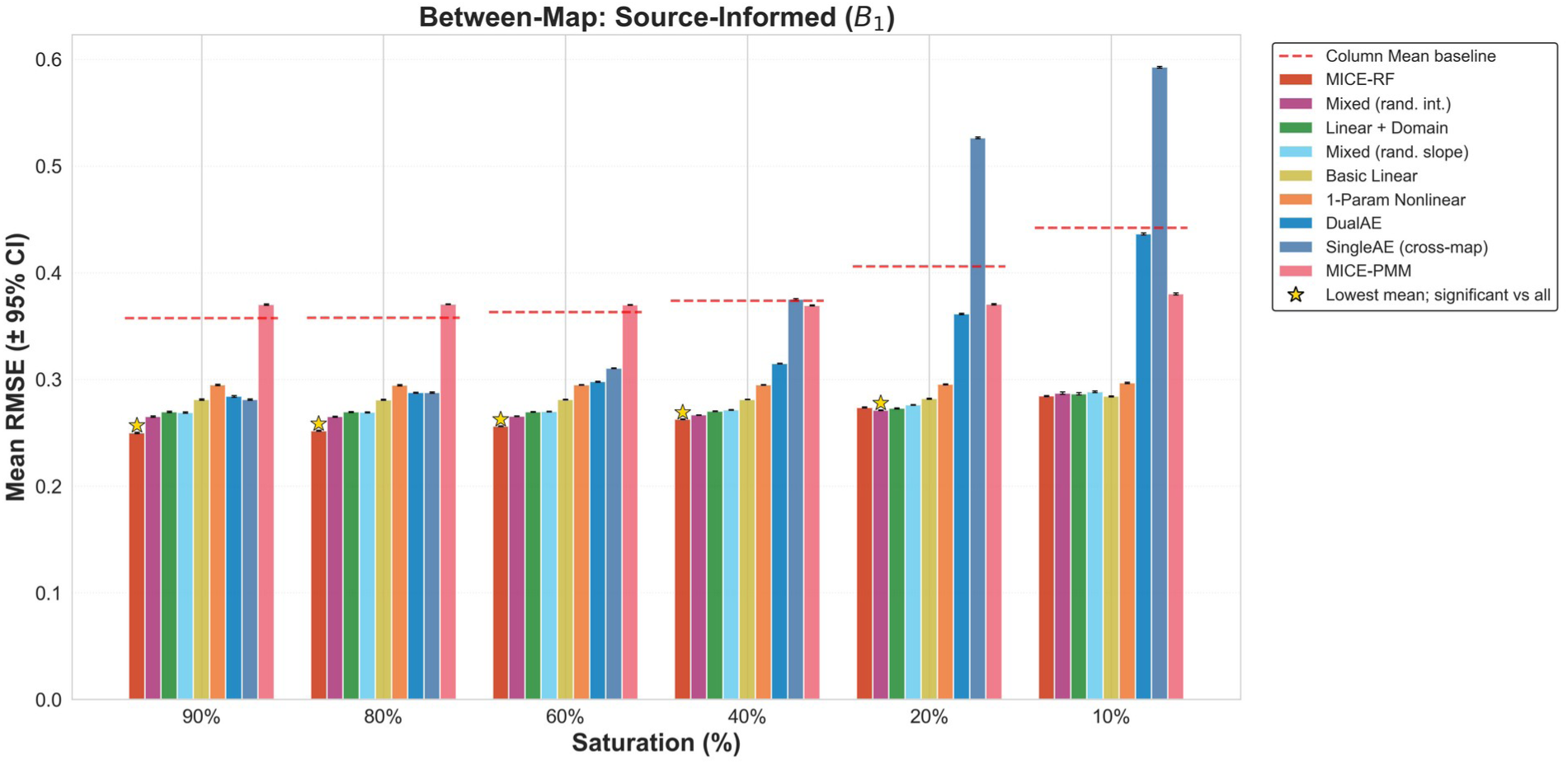
Between-map source-informed imputation accuracy. Nine source-informed methods are compared for source-informed (B_1_) held-out target values, for which the same-variant source score is available. The dashed red line shows the target-only Column Mean reference evaluated on the same B_1_ held-out values. Bars show mean RMSE on held-out values across 50 random Monte Carlo splits. Error bars show 95% confidence intervals for the mean across splits. A gold star marks the displayed method with the lowest mean only when its two-sided unpaired Mann–Whitney U test against every other displayed method remains significant after Bonferroni correction for that task and saturation. Column Mean was included in these comparisons.

Linear and mixed-effects models proved competitive. In particular, the Mixed (rand. int.) model, which uses amino acid identity as a random intercept and allows the fixed effects to vary by protein domain, achieved the second-lowest RMSE at 90% saturation (0.265), with the Linear + Domain model performing comparably (0.270). More strikingly, these models exhibited remarkable stability across saturation levels: the Basic Linear model’s RMSE increased (worsened) from 0.281 at 90% saturation to only 0.284 (only ∼1% higher) at 10% saturation. These models have few parameters to estimate, which may explain why they can be well-specified even when training data become very sparse. Indeed, at 10% saturation the Basic Linear model achieved essentially the same RMSE as MICE-RF (0.284).

Autoencoder-based methods were competitive at high saturation but degraded sharply at lower saturation. SingleAE (cross-map) achieved RMSE 0.281 at 90% saturation but deteriorated to 0.593 at 10% saturation, an approximately 111% increase, making it the worst-performing between-map method at low saturation. DualAE degraded less severely (0.284 to 0.437) but still showed far greater sensitivity to data scarcity than linear or MICE-based methods. MICE-PMM was the worst-performing between-map method at 60% saturation and above (RMSE 0.370 at 90% saturation), with SingleAE (cross-map) overtaking it as the worst performer at 40% saturation and below (0.527 and 0.593 at 20% and 10% saturation vs. MICE-PMM 0.370 and 0.380). These performance patterns reflect the higher parameter count of autoencoders, which can result in greater overfitting at lower saturation.

### Between-map missing-source (B₀) imputation: The hardest task

We next evaluated the B_0_ sub-task, in which a source map is available but the variant is missing in both source and target. Only four of our between-map methods can handle this scenario: SingleAE (cross-map), DualAE, MICE-PMM, and MICE-RF. The target-only Column Mean reference was also evaluated on the same B_0_ held-out values. Linear and mixed-effects models require a source observation as input and therefore cannot make predictions when the source value is also missing.

Missing-source imputation was substantially harder than source-informed imputation. The magnitude of the gap varied widely across models: MICE-RF showed the largest increase in error (B_1_ 0.250 vs. B_0_ 0.482 at 90% saturation, a 93% increase), while DualAE showed a more modest increase in error (B_1_ 0.298 vs. B_0_ 0.355 at 60% saturation, a 19% increase), and the gap narrowed at low saturation as B_1_ performance also degraded. This is consistent with our assertion that, without a direct source observation, models must infer a target score from patterns learned across other variants, making predictions less accurate.

DualAE achieved the best (lowest-error) B_0_ performance across most saturation levels, from 90% through 20% (RMSE 0.386 at 90% saturation, 0.364 at 40% saturation, 0.404 at 20% saturation). Its dual-decoder architecture — one decoder for reconstructing the source and one for predicting the target — enables the model to learn a latent representation that captures cross-context structure even in the absence of direct source observations. This architecture encourages the model to learn a hidden representation that can be used to both reconstruct and predict maps, making it a good candidate for tackling the B_0_ case, at the cost of poorer performance at the very lowest saturation. Indeed, when saturation fell to 10%, the target-only Column Mean reference achieved the lowest observed RMSE (0.450), followed by MICE-RF (0.456) and DualAE (0.468). MICE-RF remained the best-performing between-map method at this saturation, although it did not outperform Column Mean. SingleAE (cross-map) degraded the most severely at low saturation (0.401 at 90% saturation to 0.606 at 10% saturation), and was the worst-performing B_0_ method at 10% and 20% saturation. MICE-PMM was the worst performer at relatively high saturation (0.583 at 90% saturation). The complete missing-source comparison is shown in Figure 4 and Table 1.

**Figure 4.**
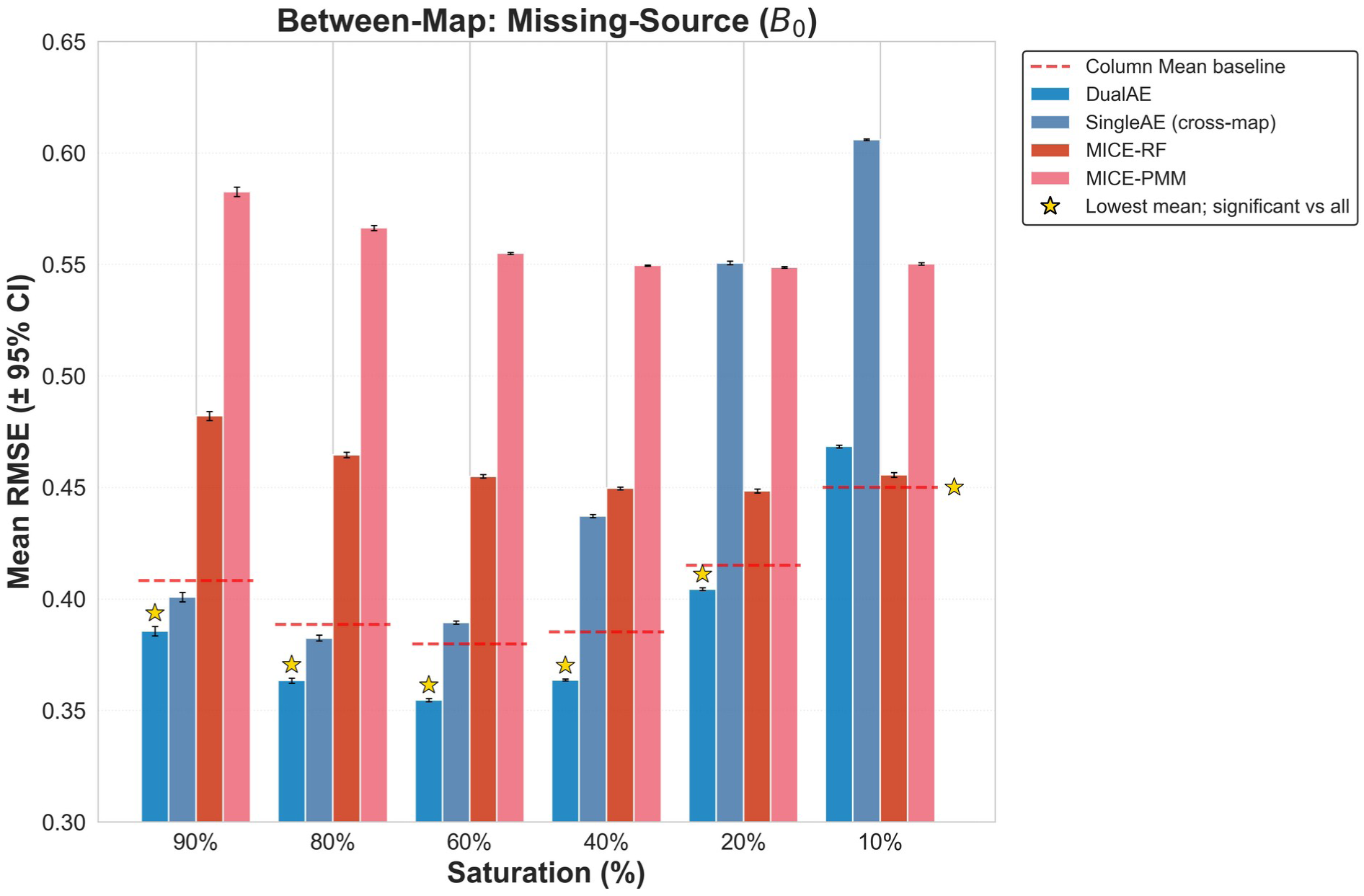
Between-map missing-source imputation accuracy. SingleAE (cross-map), DualAE, MICE-PMM, and MICE-RF are compared for missing-source (B_0_) held-out target values, for which the same-variant source score is unavailable. The dashed red line shows the target-only Column Mean reference evaluated on the same B_0_ held-out values. Bars show mean RMSE on held-out values across 50 random Monte Carlo splits. Error bars show 95% confidence intervals for the mean across splits. A gold star marks the displayed method with the lowest mean only when its two-sided unpaired Mann–Whitney U test against every other displayed method remains significant after Bonferroni correction for that task and saturation. Column Mean was included in these comparisons.

### Between-map source-informed imputation at extreme target sparsity

The B_1_ benchmark above downsampled all eight maps uniformly at random to have the same saturation, which limits how far the analysis can be pushed (at 10% saturation, only about 1% of entries are co-observed in both maps for training). To probe B_1_ behavior at saturations lower than 10%, we performed an analysis with an unmasked source map in which all originally observed source scores were retained while only the target map was subsampled. As nearly all values were present in the source map, this largely eliminated the B_0_ scenario, allowing the imputation task to remain tractable even at very low saturation levels. Here we reduced saturation down to 1% and 0.1%, at which approximately 120 and 12 target-map values, respectively, were observed per split. Results are shown in Figure 5 and Table S1.

**Figure 5.**
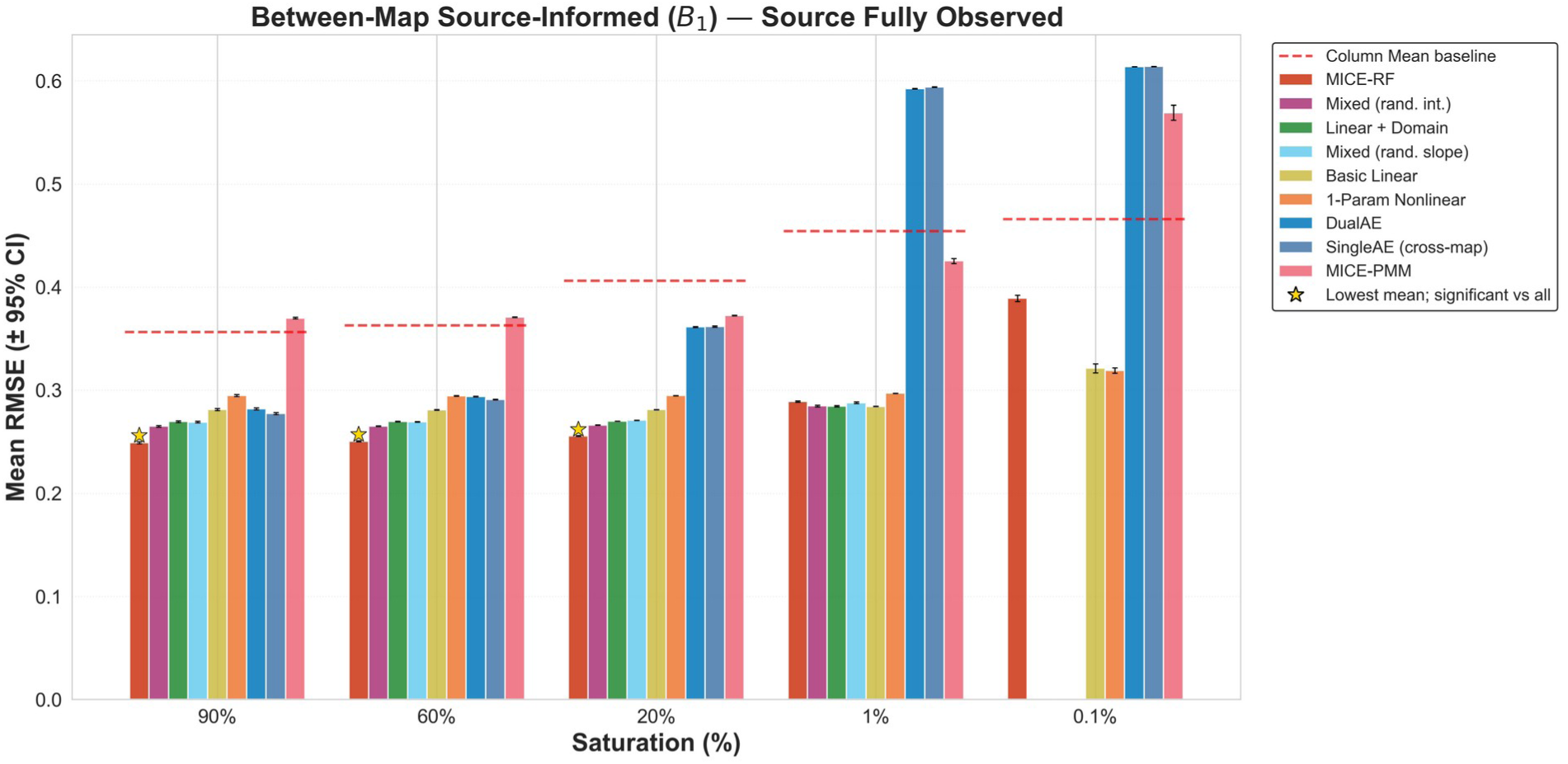
Between-map source-informed imputation at extreme target sparsity. All originally observed source-map scores are retained while target-map saturation decreases from 90% to 0.1%, largely isolating source-informed (B_1_) prediction. Naturally absent source values remain absent, and the displayed RMSE is calculated only for source-informed (B_1_) held-out values. The dashed red line shows the target-only Column Mean reference evaluated on those same held-out values. Bars show mean RMSE on held-out values across 50 random Monte Carlo splits. Error bars show 95% confidence intervals for the mean across splits. A gold star marks the displayed method with the lowest mean only when its two-sided unpaired Mann–Whitney U test against every other displayed method remains significant after Bonferroni correction for that task and saturation. Column Mean was included in these comparisons. Model–saturation combinations that did not meet the specified 95% output-completeness criterion are omitted; each comparison is restricted to the methods retained at that saturation.

At 1% target-map saturation, the linear and mixed-effects models collectively retained the strongest performance (Basic Linear 0.284 RMSE, Linear + Domain 0.284, Mixed (rand. int.) 0.285, Mixed (rand. slope) 0.288, MICE-RF 0.289), with 1-Param Nonlinear close behind (0.297). The performance of autoencoders collapsed (SingleAE (cross-map) 0.594, DualAE 0.592), as did that of MICE-PMM (0.425). At 0.1% target-map saturation, 1-Param Nonlinear achieved the lowest mean RMSE (0.319), narrowly ahead of Basic Linear (0.321); the two were not significantly different after correction. Notably, the richer regression specifications, which had been top performers at all higher saturations, did not meet the output-completeness criterion at 0.1% saturation (Linear + Domain: 43.2%; Mixed (rand. int.): 41.0%; Mixed (rand. slope): 0%) and were excluded from headline comparisons. MICE-RF, which dominated at moderate saturation, did not fall quite as far in performance but still trailed with an RMSE of 0.389, reflecting the difficulty of estimating a forest from ∼12 target values even with an unmasked source map and auxiliary covariates.

Two practical conclusions can be drawn. First, when an unmasked source map is paired with an almost-empty target map, the simplest parametric regressions recover near-best fidelity, a regime in which no autoencoder and no flexible tree model can compete. Second, model complexity pays off across most of the saturation range but becomes a liability at extreme sparsity. Every model with per-domain or per-amino-acid random effects failed the output-completeness criterion at 0.1% target-map saturation, while the simplest parametric fits remained near RMSE 0.32.

### Post hoc task-routed oracle at low saturation

While linear and mixed-effects models offer strong, stable accuracy for the B_1_ task where a direct source observation is available, they cannot make predictions when the source value is also missing (B_0_). As saturation decreases, the proportion of missing-source variants increases and the coverage (fraction of variants for which the method offers a prediction) afforded by B_1_-only methods drops dramatically; Figure 6 illustrates this shift for the ordered wt12 → av12 pair. When comparing the performance for B_1_ imputation, the best-performing method often changed dramatically depending on saturation (Figure 7). Linear models showed near-flat performance across a wide range of saturation levels for the B_1_ task, while autoencoders exhibited poor accuracy at low saturation but improved steeply as saturation increased. For B_1_ predictions, MICE-RF occupied an intermediate position between these extremes.

**Figure 6.**
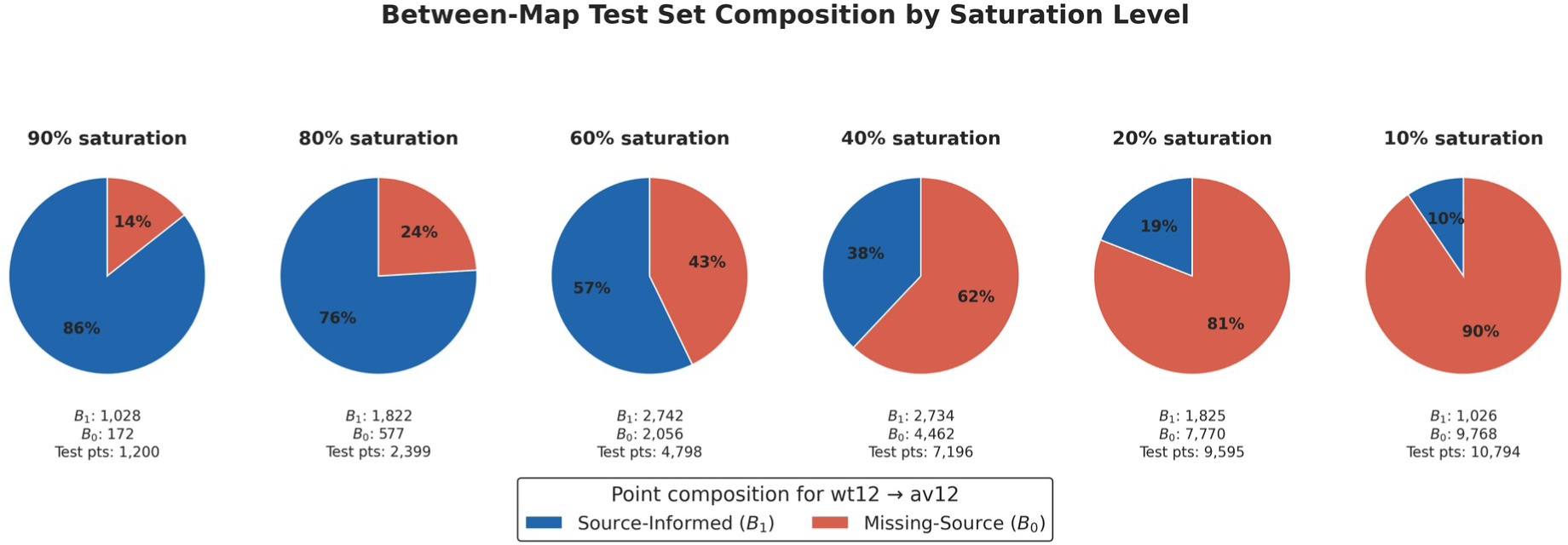
Source-informed and missing-source composition for one ordered map pair. For the ordered source-to-target pair wt12 → av12, the pie charts divide held-out values used for between-map evaluation into source-informed (B_1_) and missing-source (B_0_) values at each saturation. Counts are averaged across 50 random Monte Carlo splits and rounded to the nearest whole value. As saturation decreases, a larger share of these held-out values lack the same-variant source score. The exact proportions apply only to this ordered map pair.

**Figure 7.**
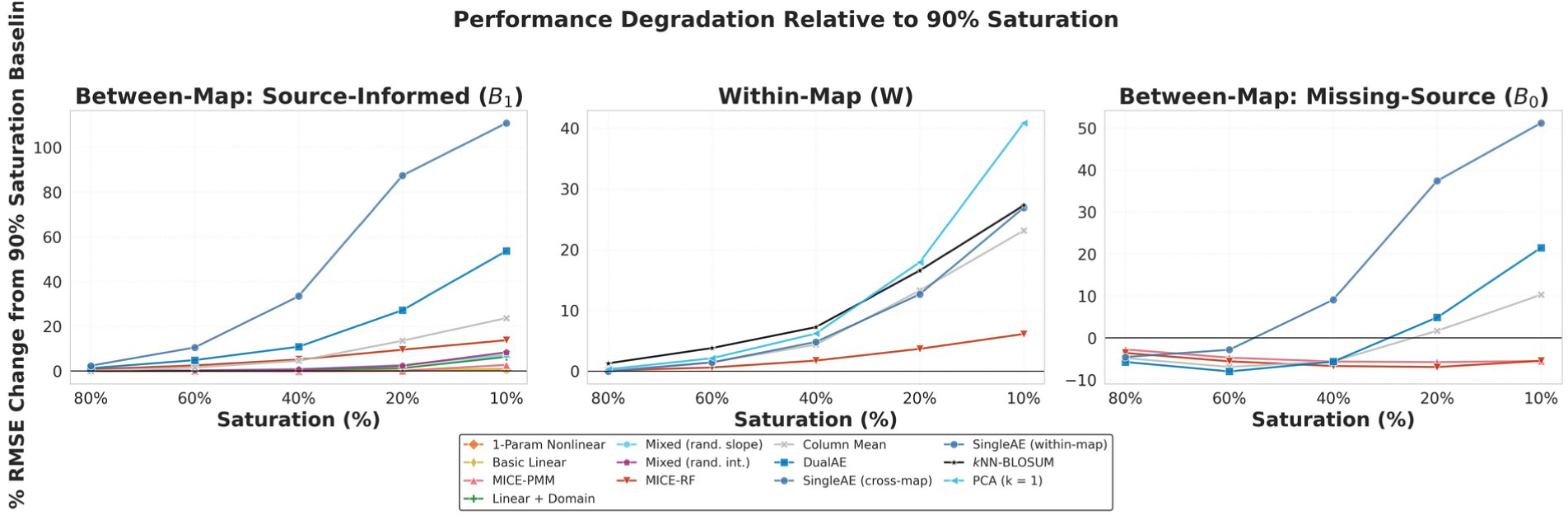
Relative change in RMSE from each model’s 90%-saturation value. For source-informed (B_1_), within-map (W), and missing-source (B_0_) evaluation, each curve shows percent change in mean RMSE on held-out values across 50 random Monte Carlo splits relative to that method’s own 90%-saturation mean. These relative changes show sensitivity to data removal and do not identify which method has the lowest RMSE. SingleAE denotes SingleAE (within-map) in the W panel and SingleAE (cross-map) in the B_1_ and B_0_ panels.

These issues motivate a “task-routed hybrid” approach: when the same-variant source score is available, the B_1_ method that generally performs best at this saturation level is used; when the same-variant source score is unavailable, the best B_0_ method at this saturation level is used. In a post hoc oracle calculation, the resulting routed summary RMSE was nominally lower than that of any evaluated single model capable of both B_1_ and B_0_ prediction across saturation levels (Figure 8). This descriptive calculation combined the selected task-level mean RMSE values and did not evaluate a trained hybrid on independent splits.

**Figure 8.**
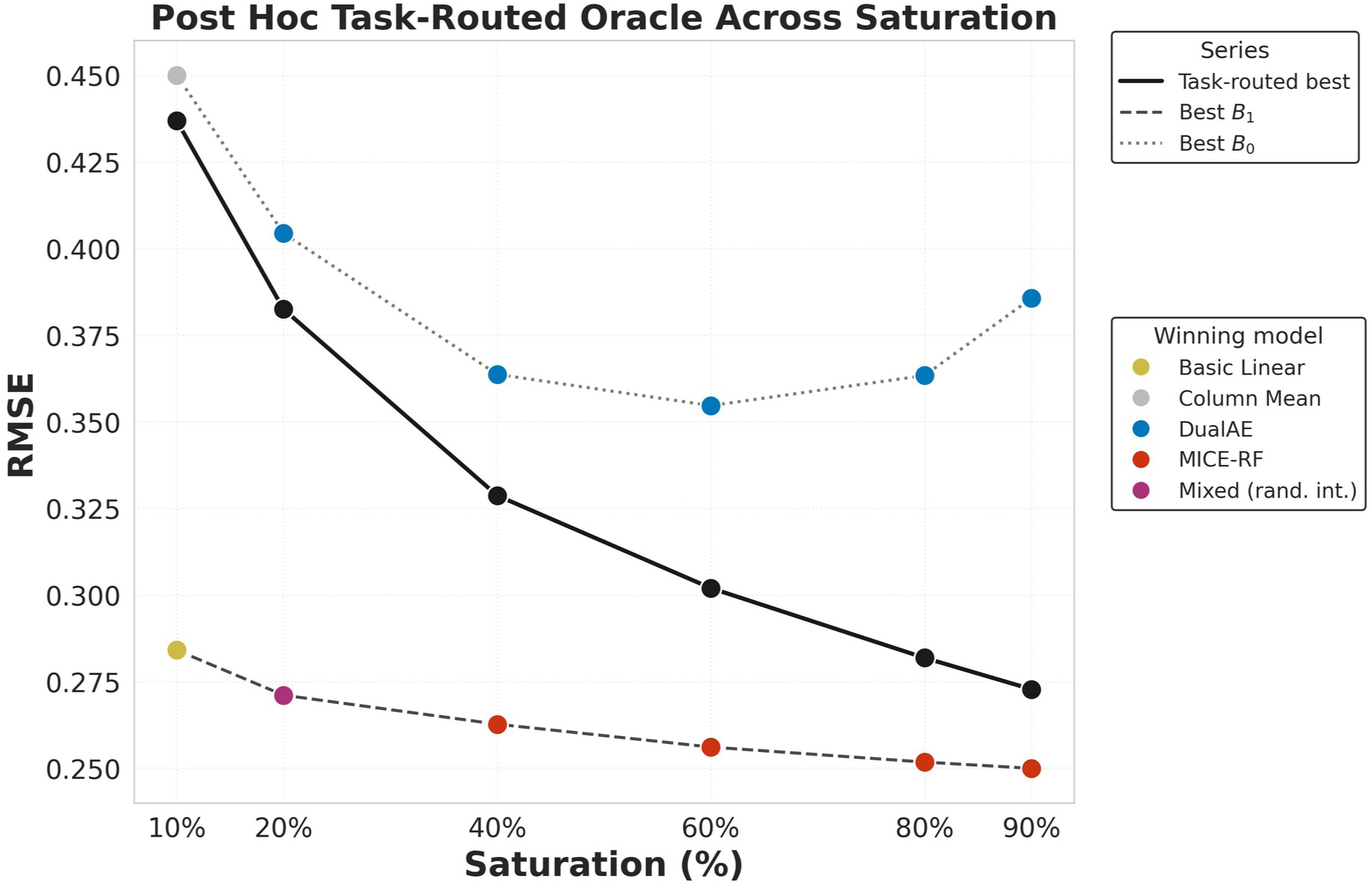
Performance of a post hoc task-routed oracle across saturation. At each saturation, the post hoc task-routed oracle separately selects the method with the lowest mean RMSE on held-out values across 50 random Monte Carlo splits for source-informed (B_1_) and missing-source (B_0_) evaluation. Selection does not require a statistically significant difference. The combined point weights the squared mean RMSE for each task by its mean number of held-out values before taking the square root. Selection included the target-only Column Mean reference. Marker color identifies the selected method for each task-specific point; black markers show the combined value. This is a descriptive summary of the methods evaluated in this study. It is not a fitted hybrid and was not evaluated on an independent set of splits. No uncertainty bars are shown because the same split results were used both to select the methods and to display their combined value.

### Handling highly position-specific context dependence

The RMSE approach summarizes overall reconstruction fidelity but can miss position-specific patterns. For example, Weile et al. identified MTHFR p.Trp165 as a residue for which non-aromatic substitutions were poorly tolerated at low folinate levels but generally tolerated at high folinate levels. Weile et al. proposed that Trp165 in a disordered loop helps retain flavin adenine dinucleotide (FAD) at the active site through aromatic π-stacking with its flavin group; folinate-dependent map effects, molecular-dynamics simulations, and enzyme assays were consistent with impaired FAD binding by non-aromatic substitutions (Weile et al. 2021). Here we performed B_1_ imputation between the highest and lowest folinate concentrations in the wild-type background. Examining RMSE for all 20 non-wild-type Trp165 outcomes (19 amino acid substitutions and one nonsense (Ter) score) across 50 randomized splits, we found autoencoder methods to have the lowest observed RMSE at Trp165 among all nine evaluated source-informed between-map methods in all 12 combinations of direction of inference and saturation level: DualAE ranked first in nine of these combinations and SingleAE ranked first in three (Table S5). The best-performing autoencoder had lower RMSE than Column Mean in nine of the 12 direction and saturation combinations. We note that Column Mean was the best method at low (40%, 20%, and 10%) saturation when imputing the low-folinate (wt12) map, reflecting the challenge of cross-context imputation in the presence of large context-dependent effects that are specific to a single residue position.

To compare imputation-side error with reported target measurement error, we calculated 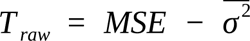 where 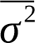 is the mean of the squared reported target standard errors (SEs) over the same held-out predictions. Assuming unbiased imputation and the measurement-error assumptions described in the Supplemental Methods, *T_raw_* represents prediction variance for a fixed mask. This prediction variance is distinct from the observed variation across the 50 randomized masks, which was summarized separately.

## Discussion

Towards extending the reach of context-specific variant effect maps by filling in missing data from sub-saturation experimental measurements, we assessed imputation methods across a range of saturation levels. Although several previous studies — for example (Weile et al. 2017; Wu et al. 2019) — have addressed within-map imputation, here we defined the between-map (B) imputation problem, in which a source map informs predictions for a target map collected under a different context. We further distinguished the source-informed (B_1_) and missing-source (B_0_) sub-tasks, because they admit different types of imputation models.

Benchmarking W imputation revealed that, from 90% to 40% saturation, PCA with one component performed best. From 20% to 10% saturation, MICE-RF (which had not, to our knowledge, been previously applied to W imputation) performed best, while kNN-BLOSUM, a within-position heuristic based on BLOSUM100, was competitive with MICE-RF.

For between-map imputation, in benchmarking B_0_ imputation, we examined DualAE, SingleAE, MICE-RF, and MICE-PMM, and found DualAE to perform best at 20% and higher saturation, whereas at 10%, Column Mean was lowest overall and MICE-RF was best among between-map methods.

In benchmarking B_1_ imputation, we found that MICE-RF achieved the lowest RMSE from 90% to 40% saturation, while the random-intercept mixed-effects model was best at 20%, and Basic Linear was nominally best and statistically indistinguishable from MICE-RF at 10% saturation. The linear and mixed-effects models provided competitive accuracy that was remarkably stable across saturation levels.

We were also able to evaluate B_1_ imputation performance at target-map saturation levels down to 0.1% by keeping the source map unmasked, with all originally observed source scores retained. At 1% target saturation, Basic Linear had the lowest mean RMSE and was statistically indistinguishable from Linear + Domain and Mixed (rand. int.); at 0.1%, 1-Param Nonlinear and Basic Linear were statistically indistinguishable.

### Why is imputation important?

Although current variant effect predictors offer near-universal coverage of missense substitutions and achieve high accuracy on benchmark sets of pathogenic and benign variants (Cheng et al. 2023; Pejaver et al. 2022; Bergquist et al. 2025), three considerations motivate continued investment in scaling functional assays through imputation. First, current VEPs produce a single static score per variant and have no explicit mechanism for incorporating the effects of environmental or genetic context on variant impacts. Indeed, as illustrated by the dependence of many MTHFR variants on the p.A222V variant and the strong folinate-dependence of most p.Trp165 substitutions, modulation of variant effects by both genetic and environmental contexts can be important (Weile et al. 2021). Training context-aware predictors would in principle close that gap, but there are few large, high-quality experimental datasets across relevant context combinations to use as training data. Imputation could help complete more context-aware datasets.

Second, most computational predictors aim to infer pathogenicity directly rather than infer impacts on protein subfunctions (stability, localization, catalytic activity, regulatory binding). For genes where pathogenicity arises from failure of a specific subfunction, an assay of that subfunction (and values imputed from it) may speak more directly to clinical pathomechanism of a variant than inferring pathogenicity more broadly (Wu et al. 2021).

Third, scaling functional measurements through cross-context imputation is a way of deriving more evidence of a type afforded high weight in current ACMG/AMP guidelines (Richards et al. 2015). Although several recent studies have supported the idea that computational predictors can provide strong evidence, in some cases approaching the discriminative power of functional assays (Frazer et al. 2021; Pejaver et al. 2022; Bergquist et al. 2025), computational and functional evidence types are synergistic rather than competitive, given that current guidelines allow use of both evidence types for the same variant (Richards et al. 2015).

### Explaining the strong performance of MICE-RF

MICE with a random forest predictor (MICE-RF) was the best-performing B_1_ method across most ordinary saturation levels, achieving the lowest RMSE from 90% to 40% saturation (0.250 at 90% saturation; 0.263 at 40% saturation), and was the best-performing between-map method for B_0_ at 10% saturation (0.456), although the target-only Column Mean reference was lower (0.450). It also achieved the lowest W RMSE at 20% and 10% saturation. One possible explanation is that MICE-RF can accommodate nonlinear, context-dependent relationships. Mutational effects can be nonlinear and context-dependent: changes in genetic background can alter abundance and binding, while allosteric coupling can propagate local perturbations to distant functional sites (Faure et al. 2022). These dependencies are likely to be especially pronounced at catalytic residues, binding interfaces, and allosteric nodes, where local perturbations propagate through long-range structural couplings and the effect of a substitution can flip from benign to deleterious (or vice versa) under a sufficiently large context shift. Therefore, it seems likely that MICE-RF performs strongly because MICE-RF, given sufficient training data, is able to learn the kinds of complicated nonlinear relationships that are intrinsic to the underlying biology.

### Strengths and limitations of the other imputation families

Linear and mixed-effects models require a source observation as input, and so were restricted to B_1_. However, for B_1_ imputation, linear and mixed-effects models were the most stable across saturation levels — the Basic Linear model’s RMSE was nearly invariant from high to extreme saturation, and at 10% saturation Basic Linear was nominally best and statistically indistinguishable from MICE-RF. The stability of linear and mixed-effects models became decisive in the asymmetric-saturation regime, where no additional source-map values were masked and the target map was almost empty. At 1% target-map saturation the simple linear and mixed-effects models collectively provided the lowest RMSEs, while every autoencoder and MICE-PMM method performed poorly (RMSE > 0.4). At 0.1% target-map saturation, with only ∼12 observed target values per split, the Basic Linear and 1-Param Nonlinear fits were the only methods that retained mean RMSE near 0.32, while the richer regression specifications did not meet the output-completeness criterion (see Results “Between-map source-informed imputation at extreme target sparsity”). This is the most resource-efficient regime a researcher would face when extending a densely measured reference map into a newly collected sparse context. Linear and mixed-effects models also have the advantage of being interpretable: a target-vs-source slope is a fitness transfer function between contexts, a domain-interaction term is a domain-specific deviation, and modelers can encode biologically meaningful structure into the design matrix. Their main weakness is that they cannot impute B_0_.

The focused Trp165 analysis helps explain why the model rankings differed across this context shift. The linear and mixed models preserved much of the relative ordering among substitutions, but their global source-to-target relationships did not reproduce the especially large shift in scores between the low- and high-folinate maps at Trp165. Autoencoders generally placed predictions closer to the overall target-map score level, although SingleAE predictions increasingly contracted toward the wild-type score of 1 at low saturation. This helped when predicting the generally high wt200 scores but hurt when predicting the lower wt12 scores, showing that lower RMSE did not always mean that the substitution-specific folinate response had been recovered. This phenomenon is exemplified by the fact that Column Mean remained a competitive approach at low saturation and was the best-performing method in the high-to-low (wt200 → wt12) folinate direction at 10% saturation despite not taking any cross-context information into account.

More parameter-rich autoencoders delivered strong B_1_ accuracy when training data were plentiful, but showed rapid performance degradation with lower saturation (e.g., the error on SingleAE (cross-map) imputation rose from RMSE 0.281 at 90% saturation to RMSE 0.593 at 10% saturation, the worst B_1_ performance of any method at 10% saturation). These methods are also largely opaque, making it hard to inject biological priors or to diagnose an individual prediction. Their compensating advantage is coverage: both SingleAE (cross-map) and DualAE can predict B_0_, and DualAE in particular delivered the best B_0_ accuracy across most of the saturation range. This matters more than it might initially seem, because the composition of the between-map test set shifts dramatically toward B_0_ as saturation decreases; Figure 6 illustrates this shift for the ordered wt12 → av12 pair.

MICE-PMM was the weakest between-map method we tested at high-to-moderate saturation (RMSE 0.370 at 90% saturation), but was included as the standard reference implementation of chained-equations imputation.

### Proposing a new pathogenicity evidence type for ‘purely experimental’ functional imputation

The case for imputation rests on more than the question of which model wins which benchmark. If imputation is to be useful for clinical variant classification, then clinical guidelines will need to recognize the special nature of evidence imputed from experimental functional data. Current ACMG/AMP guidelines (Richards et al. 2015) place computational evidence (PP3, BP4) and experimental functional evidence (PS3, BS3) in distinct categories, potentially each with its own calibration framework (Pejaver et al. 2022; Bergquist et al. 2025; Brnich et al. 2020). Importantly, values imputed from experimental evidence do not fit cleanly into either category. Although these values are derived computationally, their primary input is experimentally measured functional data rather than the sequence, evolutionary, and structural features used by many VEPs (Gao et al. 2023; Jagota et al. 2023; Gerasimavicius et al. 2025). This distinction may be especially relevant when an assay measures a disease-relevant function that is not adequately represented by the features used by conventional VEPs. We therefore propose that the evidentiary weight assigned to an imputed functional score should reflect both the validated ability of the underlying assay to distinguish independently classified benign and pathogenic control variants and the predictive reliability of the imputation procedure. The former can be quantified when sufficient validation data are available (Brnich et al. 2020). The latter should be evaluated empirically using held-out or prospectively measured variants. Some probabilistic genotype–phenotype reconstruction methods also provide model-based uncertainty estimates for individual predictions of unmeasured variant phenotypes (Zhou et al. 2022), but such estimates would themselves require empirical calibration before they could inform evidence strength. A clinical framework that explicitly accounts for both components would be more appropriate than one that bins imputed values into the existing computational category.

A dedicated evidence code would also clarify a question the current framework leaves implicit. How dependent are the various types of evidence on one another? Such a code would be conceptually adjacent to but distinct from PM5, which is awarded when another known pathogenic substitution exists at the same position. The current framework defines PP3/BP4, PS3/BS3, and PM5 as distinct criteria but does not explicitly model their error dependence. Some clinical geneticists treat computational and functional evidence as fully dependent. Two considerations motivate this rationale: (1) both approaches assess variant impact on protein-level function; and (2) to the extent that pathogenicity and protein dysfunction are not fully aligned, the two evidence types will have correlated errors. Imputed functional evidence sits between VEP and functional evidence, forcing the framework to make its dependence structure explicit. Working out the relevant dependence relationships among PP3/BP4, PS3/BS3, PM5, and a putative experimentally imputed evidence type has the potential to improve the accuracy of clinical variant classification.

### Limitations and future directions

This study has several limitations. First, we evaluated imputation performance using RMSE on uniformly random held-out variants, which is the appropriate approach when the experimental sub-saturation strategy is also uniformly random. However, a variety of experimental sub-saturation strategies are possible, including: saturated alanine scanning, which replaces each residue in turn with alanine and yields ∼5% saturation (Cunningham and Wells 1989); histidine- and asparagine-only scanning, which yields ∼11% saturation (Gray et al. 2017); and single-nucleotide-only mutagenesis, which cannot measure all possible amino acid substitutions (Findlay et al. 2018). Although each sub-saturation strategy induces a different test-set composition and could in principle be evaluated against the present saturated MTHFR data, we did not attempt that here. The MTHFR dataset offered an ideal testbed, given the availability of MAVEs measuring dependence of variant impacts on both environmental and genetic contexts. Future work could assess the generality of our findings for a wider range of genes and contexts. Future work might also weight evaluation toward variants of higher clinical interest rather than treating all held-out variants equally.

Second, aggregate RMSE can obscure position-specific and directional differences in model performance, as illustrated by the Trp165 analysis.

Third, we deliberately kept the imputation pipeline free of computational VEP signals to minimize dependence between these evidence types and avoid double-counting when evidence from imputed values and VEPs is combined. Relaxing that constraint by combining imputation with current-generation VEPs would almost certainly improve point estimates, but at the cost of introducing additional dependence between VEPs and imputed functional evidence. A separate direction is physics- and biology-informed models that incorporate constraints from known protein behavior (folding stability, allosteric coupling, active-site geometry); these have not yet been brought to bear on cross-context MAVE imputation and are a promising avenue.

Finally, although our framing has been about imputing missing values, imputation is also useful even where functional measurements are available. A weighted combination of measured and imputed scores can yield a refined estimate of the true functional impact whose error can be lower than either input alone when their errors are sufficiently independent and the weights are appropriately calibrated, an idea developed in earlier MAVE work (Weile et al. 2017). Gaussian-process formulations make this weighting explicit by conditioning jointly on the observed scores and on the experimental error variance attached to each one, so that noisier measurements are shrunk further toward the model prediction (Zhou et al. 2022).

Taken together, our findings provide concrete recommendations for researchers allocating finite experimental budgets across multiple contexts: 1) choose MICE-RF as the default cross-context imputer when a source map is available and saturation is moderate to high; 2) use simple linear or mixed-effects models at lower target saturation; 3) use the simplest regression models at very low saturation (and when an unmasked source map is available); 4) use DualAE when the missing-source case dominates and saturation is sufficient to support an autoencoder; and 5) consider treating the resulting imputed values as a distinct class of clinical evidence that warrants its own place in the framework for clinical variant interpretation.

## Supporting information

upplemental Information

Statistical Results and Supporting CSV Files

## Acknowledgments

Funding: We acknowledge funding for this work from the National Human Genome Research Institute (NHGRI) of the National Institutes of Health (NIH) Center of Excellence in Genomic Science Initiative (RM1HG010461; to F.P.R.) and NIH National Cancer Institute (NCI; R01CA313942; to R.B.). F.P.R. was also supported by the NHGRI Impact of Genomic Variation on Function Initiative UM1HG011989.

## Author contributions

Conceptualization: A.R., R.B., and F.P.R. Methodology: A.R. and R.B. Software: A.R. Validation: A.R. Formal analysis: A.R. Data curation: A.R. and V.S. Visualization: A.R. and W.v.L. Resources: F.P.R. Funding acquisition: R.B. and F.P.R. Project administration: A.R. and F.P.R. Supervision: R.B. and F.P.R. Writing (original draft): A.R. Writing (review & editing): A.R., R.B., and F.P.R.

## Data and code availability

Intermediate split files and derived analysis outputs can be regenerated using the scripts and documented parameters in the repository. All source data and analysis code used in this study are available on GitHub (commit d279e755): https://github.com/aliras11/MAVECrossContext_Imputation.

## Declaration of generative AI and AI-assisted technologies

During the preparation of this work, the authors used OpenAI Codex and Anthropic Claude to assist with revising and expanding author-developed research code; checking file paths; preparing and managing job-submission scripts; generating reproducible figure-production code from detailed author specifications; formatting citations; checking grammar; cross-checking reported numerical claims against the underlying data; and performing submission-readiness checks. The initial implementations, scientific questions, analytical decisions, and interpretation of results were developed by the authors. The authors reviewed and tested all AI-assisted code, verified the analyses and figures against the underlying data, reviewed and edited the manuscript, and take full responsibility for the content of this work.

## References

Timothy Bergquist, Sarah L Stenton, Emily A W Nadeau, Alicia B Byrne, Marc S Greenblatt, Steven M Harrison, Sean V Tavtigian, Anne O’Donnell-Luria, Leslie G Biesecker, Predrag Radivojac, Steven E Brenner, Vikas Pejaver, and ClinGen Sequence Variant Interpretation Working Group. Calibration of additional computational tools expands ClinGen recommendation options for variant classification with PP3/BP4 criteria. Genet. Med., 27 (6): 101402, June 2025. doi:10.1016/j.gim.2025.101402.

Linnea K M Blomgren, Melanie Huber, Sabrina R Mackinnon, Céline Bürer, Arnaud Baslé, Wyatt W Yue, D Sean Froese, and Thomas J McCorvie. Dynamic inter-domain transformations mediate the allosteric regulation of human 5,10-methylenetetrahydrofolate reductase. Nat. Commun., 15 (1): 3248, April 2024. doi:10.1038/s41467-024-47174-y.

Sarah E Brnich, Ahmad N Abou Tayoun, Fergus J Couch, et al., on behalf of the Clinical Genome Resource Sequence Variant Interpretation Working Group. Recommendations for application of the functional evidence PS3/BS3 criterion using the ACMG/AMP sequence variant interpretation framework. Genome Med., 12 (1): 3, 2020. doi:10.1186/s13073-019-0690-2.

Jun Cheng, Guido Novati, Joshua Pan, et al. Accurate proteome-wide missense variant effect prediction with AlphaMissense. Science, 381 (6664): eadg7492, September 2023. doi:10.1126/science.adg7492.

B C Cunningham and J A Wells. High-resolution epitope mapping of hGH-receptor interactions by alanine-scanning mutagenesis. Science, 244 (4908): 1081–1085, June 1989. doi:10.1126/science.2471267.

Andre J Faure, Júlia Domingo, Jörn M Schmiedel, Cristina Hidalgo-Carcedo, Guillaume Diss, and Ben Lehner. Mapping the energetic and allosteric landscapes of protein binding domains. Nature, 604 (7904): 175–183, April 2022. doi:10.1038/s41586-022-04586-4.

Gregory M Findlay, Riza M Daza, Beth Martin, Melissa D Zhang, Anh P Leith, Molly Gasperini, Joseph D Janizek, Xingfan Huang, Lea M Starita, and Jay Shendure. Accurate classification of BRCA1 variants with saturation genome editing. Nature, 562 (7726): 217–222, October 2018. doi:10.1038/s41586-018-0461-z.

Douglas M Fowler and Heidi L Rehm. Will variants of uncertain significance still exist in 2030? Am. J. Hum. Genet., 111 (1): 5–10, January 2024. doi:10.1016/j.ajhg.2023.11.005.

Douglas M Fowler, David J Adams, Anna L Gloyn, William C Hahn, Debora S Marks, Lara A Muffley, James T Neal, Frederick P Roth, Alan F Rubin, Lea M Starita, and Matthew E Hurles. An atlas of variant effects to understand the genome at nucleotide resolution. Genome Biol., 24 (1): 147, July 2023. doi:10.1186/s13059-023-02986-x.

Jonathan Frazer, Pascal Notin, Mafalda Dias, et al. Disease variant prediction with deep generative models of evolutionary data. Nature, 599 (7883): 91–95, November 2021. doi:10.1038/s41586-021-04043-8.

Hong Gao, Tobias Hamp, Jeffrey Ede, Joshua G Schraiber, Jeremy McRae, Moriel Singer-Berk, Yanshen Yang, Anastasia S D Dietrich, Petko P Fiziev, Lukas F K Kuderna, Laksshman Sundaram, Yibing Wu, Aashish Adhikari, Yair Field, Chen Chen, Serafim Batzoglou, Francois Aguet, Gabrielle Lemire, Rebecca Reimers, Daniel Balick, Mareike C Janiak, Martin Kuhlwilm, Joseph D Orkin, Shivakumara Manu, Alejandro Valenzuela, Juraj Bergman, Marjolaine Rousselle, Felipe Ennes Silva, Lidia Agueda, Julie Blanc, Marta Gut, Dorien de Vries, Ian Goodhead, R Alan Harris, Muthuswamy Raveendran, Axel Jensen, Idriss S Chuma, Julie E Horvath, Christina Hvilsom, David Juan, Peter Frandsen, Fabiano R de Melo, Fabrício Bertuol, Hazel Byrne, Iracilda Sampaio, Izeni Farias, João Valsecchi do Amaral, Mariluce Messias, Maria N F da Silva, Mihir Trivedi, Rogerio Rossi, Tomas Hrbek, Nicole Andriaholinirina, Clément J Rabarivola, Alphonse Zaramody, Clifford J Jolly, Jane Phillips-Conroy, Gregory Wilkerson, Christian Abee, Joe H Simmons, Eduardo Fernandez-Duque, Sree Kanthaswamy, Fekadu Shiferaw, Dongdong Wu, Long Zhou, Yong Shao, Guojie Zhang, Julius D Keyyu, Sascha Knauf, Minh D Le, Esther Lizano, Stefan Merker, Arcadi Navarro, Thomas Bataillon, Tilo Nadler, Chiea Chuen Khor, Jessica Lee, Patrick Tan, Weng Khong Lim, Andrew C Kitchener, Dietmar Zinner, Ivo Gut, Amanda Melin, Katerina Guschanski, Mikkel Heide Schierup, Robin M D Beck, Govindhaswamy Umapathy, Christian Roos, Jean P Boubli, Monkol Lek, Shamil Sunyaev, Anne O’Donnell-Luria, Heidi L Rehm, Jinbo Xu, Jeffrey Rogers, Tomas Marques-Bonet, and Kyle Kai-How Farh. The landscape of tolerated genetic variation in humans and primates. Science, 380 (6648): eabn8153, June 2023. doi:10.1126/science.abn8197.

Marinella Gebbia, Daniel Zimmerman, Rosanna Jiang, Maria Nguyen, Jochen Weile, Roujia Li, Michelle Gavac, Nishka Kishore, Song Sun, Rick A Boonen, Rayna Hamilton, Jennifer N Dines, Alexander Wahl, Jason Reuter, Britt Johnson, Douglas M Fowler, Fergus J Couch, Haico van Attikum, and Frederick P Roth. A missense variant effect map for the human tumor-suppressor protein CHK2. Am. J. Hum. Genet., 111 (12): 2675–2692, December 2024. doi:10.1016/j.ajhg.2024.10.013.

Lukas Gerasimavicius, Sarah A Teichmann, and Joseph A Marsh. Leveraging protein structural information to improve variant effect prediction. Curr. Opin. Struct. Biol., 92: 103023, June 2025. doi:10.1016/j.sbi.2025.103023.

Vanessa E Gray, Ronald J Hause, and Douglas M Fowler. Analysis of large-scale mutagenesis data to assess the impact of single amino acid substitutions. Genetics, 207 (1): 53–61, September 2017. doi:10.1534/genetics.117.300064.

Dominik G Grimm, Chloé-Agathe Azencott, Fabian Aicheler, Udo Gieraths, Daniel G MacArthur, Kaitlin E Samocha, David N Cooper, Peter D Stenson, Mark J Daly, Jordan W Smoller, Laramie E Duncan, and Karsten M Borgwardt. The evaluation of tools used to predict the impact of missense variants is hindered by two types of circularity. Hum. Mutat., 36 (5): 513–523, 2015. doi:10.1002/humu.22768.

S Henikoff and J G Henikoff. Amino acid substitution matrices from protein blocks. Proc. Natl. Acad. Sci. U. S. A., 89 (22): 10915–10919, November 1992. doi:10.1073/pnas.89.22.10915.

Martina Huemer and Matthias R Baumgartner. The clinical presentation of cobalamin-related disorders: From acquired deficiencies to inborn errors of absorption and intracellular pathways. J. Inherit. Metab. Dis., 42 (4): 686–705, July 2019. doi:10.1002/jimd.12012.

Milind Jagota, Chengzhong Ye, Carlos Albors, Ruchir Rastogi, Antoine Koehl, Nilah Ioannidis, and Yun S Song. Cross-protein transfer learning substantially improves disease variant prediction. Genome Biol., 24 (1): 182, August 2023. doi:10.1186/s13059-023-03024-6.

Ben Langmead and Steven L Salzberg. Fast gapped-read alignment with Bowtie 2. Nat. Methods, 9 (4): 357– 359, 2012. doi:10.1038/nmeth.1923.

Siaw-Cheok Liew and Esha Das Gupta. Methylenetetrahydrofolate reductase (MTHFR) C677T polymorphism: epidemiology, metabolism and the associated diseases. Eur. J. Med. Genet., 58 (1): 1–10, January 2015. doi:10.1016/j.ejmg.2014.10.004.

Benjamin J Livesey and Joseph A Marsh. Variant effect predictor correlation with functional assays is reflective of clinical classification performance. Genome Biol., 26 (1): 104, April 2025. doi:10.1186/s13059-025-03575-w.

Vikas Pejaver, Alicia B Byrne, Bing-Jian Feng, Kymberleigh A Pagel, Sean D Mooney, Rachel Karchin, Anne O’Donnell-Luria, Steven M Harrison, Sean V Tavtigian, Marc S Greenblatt, Leslie G Biesecker, Predrag Radivojac, Steven E Brenner, and ClinGen Sequence Variant Interpretation Working Group. Calibration of computational tools for missense variant pathogenicity classification and ClinGen recommendations for PP3/BP4 criteria. Am. J. Hum. Genet., 109 (12): 2163–2177, December 2022. doi:10.1016/j.ajhg.2022.10.013.

Polina V Polunina, Wolfgang Maier, and Alan F Rubin. VEFill: accurate and generalizable deep mutational scanning score imputation across protein domains. Mol. Syst. Biol., 22 (6): 979–1002, June 2026. doi:10.1038/s44320-026-00203-y.

Sue Richards, Nazneen Aziz, Sherri Bale, David Bick, Soma Das, Julie Gastier-Foster, Wayne W Grody, Madhuri Hegde, Elaine Lyon, Elaine Spector, Karl Voelkerding, and Heidi L Rehm, on behalf of the ACMG Laboratory Quality Assurance Committee. Standards and guidelines for the interpretation of sequence variants: a joint consensus recommendation of the American College of Medical Genetics and Genomics and the Association for Molecular Pathology. Genet. Med., 17 (5): 405–424, May 2015. doi:10.1038/gim.2015.30.

Paulus S Rommer, Johannes Zschocke, Brian Fowler, Manuela Födinger, Vassiliki Konstantopoulou, Dorothea Möslinger, Elisabeth Stögmann, Erhard Suess, Matthias Baumgartner, Eduard Auff, and Gere Sunder- Plassmann. Manifestations of neurological symptoms and thromboembolism in adults with MTHFR-deficiency. J. Neurol. Sci., 383: 123–127, December 2017. doi:10.1016/j.jns.2017.10.035.

Donald B Rubin. Inference and missing data. Biometrika, 63 (3): 581–592, December 1976. doi:10.1093/biomet/63.3.581.

Jay Shendure and Joshua M Akey. The origins, determinants, and consequences of human mutations. Science, 349 (6255): 1478–1483, September 2015. doi:10.1126/science.aaa9119.

Lea M Starita, Nadav Ahituv, Maitreya J Dunham, Jacob O Kitzman, Frederick P Roth, Georg Seelig, Jay Shendure, and Douglas M Fowler. Variant interpretation: Functional assays to the rescue. Am. J. Hum. Genet., 101 (3): 315–325, September 2017. doi:10.1016/j.ajhg.2017.07.014.

Daniel Tabet, Victoria Parikh, Prashant Mali, Frederick P Roth, and Melina Claussnitzer. Scalable functional assays for the interpretation of human genetic variation. Annu. Rev. Genet., 56: 441–465, November 2022. doi:10.1146/annurev-genet-072920-032107.

Daniel R Tabet, Da Kuang, Megan C Lancaster, Roujia Li, Karen Liu, Jochen Weile, Atina G Coté, Yingzhou Wu, Robert A Hegele, Dan M Roden, and Frederick P Roth. Benchmarking computational variant effect predictors by their ability to infer human traits. Genome Biol., 25 (1): 172, July 2024. doi:10.1186/s13059-024-03314-7.

Stef van Buuren and Karin Groothuis-Oudshoorn. mice: Multivariate imputation by chained equations in R. J. Stat. Softw., 45 (3): 1–67, December 2011. doi:10.18637/jss.v045.i03.

Pascal Vincent, Hugo Larochelle, Yoshua Bengio, and Pierre-Antoine Manzagol. Extracting and composing robust features with denoising autoencoders. In Proceedings of the 25th international conference on Machine learning, ICML’08, page 1096–1103, New York, NY, USA, July 2008. Association for Computing Machinery. doi:10.1145/1390156.1390294.

Jochen Weile, Song Sun, Atina G Cote, Jennifer Knapp, Marta Verby, Joseph C Mellor, Yingzhou Wu, Carles Pons, Cassandra Wong, Natascha van Lieshout, Fan Yang, Murat Tasan, Guihong Tan, Shan Yang, Douglas M Fowler, Robert Nussbaum, Jesse D Bloom, Marc Vidal, David E Hill, Patrick Aloy, and Frederick P Roth. A framework for exhaustively mapping functional missense variants. Mol. Syst. Biol., 13 (12): 957, December 2017. doi:10.15252/msb.20177908.

Jochen Weile, Nishka Kishore, Song Sun, Ranim Maaieh, Marta Verby, Roujia Li, Iosifina Fotiadou, Julia Kitaygorodsky, Yingzhou Wu, Alexander Holenstein, Céline Bürer, Linnea Blomgren, Shan Yang, Robert Nussbaum, Rima Rozen, David Watkins, Marinella Gebbia, Viktor Kozich, Michael Garton, D Sean Froese, and Frederick P Roth. Shifting landscapes of human MTHFR missense-variant effects. Am. J. Hum. Genet., 108 (7): 1283–1300, July 2021. doi:10.1016/j.ajhg.2021.05.009.

Sam Wilson. miceRanger: Multiple imputation by chained equations with random forests, September 2021. URL: https://CRAN.R-project.org/package=miceRanger. R package, CRAN release 1.5.0 (2021-09-06). doi:10.32614/CRAN.package.miceRanger.

Yingzhou Wu, Jochen Weile, Atina G Cote, Song Sun, Jennifer Knapp, Marta Verby, and Frederick P Roth. A web application and service for imputing and visualizing missense variant effect maps. Bioinformatics, 35 (17): 3191–3193, September 2019. doi:10.1093/bioinformatics/btz012.

Yingzhou Wu, Roujia Li, Song Sun, Jochen Weile, and Frederick P Roth. Improved pathogenicity prediction for rare human missense variants. Am. J. Hum. Genet., 108 (10): 1891–1906, October 2021. doi:10.1016/j.ajhg.2021.08.012.

Tian Yu, James D Fife, Vineel Bhat, Ivan Adzhubey, Richard Sherwood, and Christopher A Cassa. FUSE: Improving the estimation and imputation of variant impacts in functional screening. Cell Genom., 4 (10): 100667, October 2024. doi:10.1016/j.xgen.2024.100667.

Juannan Zhou, Mandy S Wong, Wei-Chia Chen, Adrian R Krainer, Justin B Kinney, and David M McCandlish. Higher-order epistasis and phenotypic prediction. Proc. Natl. Acad. Sci. U. S. A., 119 (39): e2204233119, September 2022. doi:10.1073/pnas.2204233119.

