## Supplementary material for "Inferring Protein Variant Impacts Across Contexts": upplemental Information: README.txt

AJHG statistical result CSVs
Prepared 2026-08-14 from the canonical 2026-08-08 submission package.

CONTENTS AND SCOPE
This archive contains the 16 latest statistical result CSVs used to support the
RMSE summaries, model comparisons, completeness checks, and significance claims
in the MTHFR MAVE imputation manuscript and supplemental information.
The CSV payloads are unchanged from the canonical statistics folder and are
byte-identical to their latest source copies. Together they contain 31,126 data rows.

PROVENANCE AND REPRODUCTION
Authoritative code: https://github.com/aliras11/MAVECrossContext_Imputation
Audited code commit: d279e7550efd38bf2c11a44981128afd57668b85
Source MTHFR maps: MaveDB urn:mavedb:00000049
Intermediate random split files, prediction files, and other derived analysis
outputs are not duplicated in this archive. They can be regenerated with the
repository scripts and documented parameters (50 splits; base seed 42).

RELATED SUPPLEMENT
Table S4 uses a separate variance-decomposition source family and is supplied as
Table_S4_variance_decomposition.xlsx; it is intentionally not one of these 16 CSVs.

FILES
- nodouble_model_rate_completeness.csv (50 rows): No-double output completeness by model and rate.
- pairwise_mwu_by_context_double_missing.csv (3,360 rows): Missing-source (B0) per-context Mann–Whitney U comparisons.
- pairwise_mwu_by_context_nodouble_regression_test.csv (11,256 rows): Asymmetric source-informed (B1) per-context comparisons.
- pairwise_mwu_by_context_regression_test.csv (15,120 rows): Regular source-informed (B1) per-context comparisons.
- pairwise_mwu_by_context_within_map.csv (480 rows): Within-map (W) per-context comparisons.
- pairwise_mwu_double_missing.csv (60 rows): Missing-source (B0) headline comparisons.
- pairwise_mwu_nodouble_regression_test.csv (201 rows): Asymmetric source-informed (B1) headline comparisons.
- pairwise_mwu_regression_test.csv (270 rows): Regular source-informed (B1) headline comparisons.
- pairwise_mwu_within_map.csv (60 rows): Within-map (W) headline comparisons.
- rmse_summary_double_missing.csv (30 rows): Missing-source (B0) RMSE summary.
- rmse_summary_nodouble_regression_test.csv (47 rows): Asymmetric source-informed (B1) RMSE summary.
- rmse_summary_regression_test.csv (60 rows): Regular source-informed (B1) RMSE summary.
- rmse_summary_within_map.csv (30 rows): Within-map (W) RMSE summary.
- vs_colmean_double_missing.csv (24 rows): Missing-source (B0) comparisons with Column Mean.
- vs_colmean_regression_test.csv (54 rows): Source-informed (B1) comparisons with Column Mean.
- vs_colmean_within_map.csv (24 rows): Within-map (W) comparisons with Column Mean.

INTEGRITY
SHA256SUMS.txt records the SHA-256 digest for each CSV in this archive.
