## Supplemental Information for "Inferring Protein Variant Impacts Across Contexts"

### Supplemental methods

#### Benchmark tasks and held-out values

We evaluated within-map (W) and between-map (B) imputation. Within-map imputation predicts a held-out target score without using measurements from a separate source-context map. Between-map imputation uses a separate source map. It is source-informed ($B_1$) when the same-variant source score is available and missing-source ($B_0$) when that score is unavailable. Saturation is the fraction of originally observed scores retained for training.

In each of 50 random Monte Carlo splits, RMSE was calculated only for target scores that were originally observed and then held out for evaluation. Entries without an original target measurement were not scored. Whether a between-map held-out value was assigned to $B_1$ or $B_0$ was determined from same-variant source-score availability in that split.

The mathematical forms and fitted parameters of the five regression models implemented in R are summarized in Table S6.

#### RMSE aggregation and statistical analysis

For each method, task, saturation, and split, we calculated the squared error for every held-out value across the applicable maps or ordered source-to-target pairs. We added those squared errors, divided by the total number of held-out values, and took the square root. Equivalently, the result for each map or pair contributes $N \times RMSE^2$ to the numerator and N to the denominator. Maps or pairs with more scored values therefore contribute more; their RMSE values are not given equal weight.

The 50 random splits yield 50 RMSE values for each method, task, and saturation combination meeting the specified 95% output-completeness criterion. Figures and summary tables report their mean and a 95% confidence interval for that mean. We compared methods using two-sided unpaired Mann–Whitney U tests on the split-level RMSE values and applied Bonferroni correction to all displayed method comparisons within each task and saturation. Separate tests for individual maps or ordered source-to-target pairs are reported in Table S3; these tests do not determine the gold stars in the main figures.

#### Task-routed oracle analysis

We summarized a post hoc task-routed oracle for between-map imputation. For each saturation and task, we first calculated one RMSE per method in each of the 50 random Monte Carlo splits across all held-out values and ordered source-to-target map pairs. We then selected the method with the lowest mean of those 50 RMSE values separately among all displayed methods, including the target-only Column Mean reference for both $B_1$ and $B_0$. Statistical significance was not required for selection. Same-variant source-score availability determines whether a held-out value belongs to $B_1$ or $B_0$.

The analysis did not fit a new hybrid model or generate new predictions. Instead, the displayed value was calculated from the selected $B_1$ and $B_0$ mean RMSE values. Each squared task-specific mean was weighted by the mean number of held-out values in that task before taking the square root. For each task and saturation, held-out values were first counted across ordered map pairs within each split and then averaged across the 50 splits. In the equations below, R_m,q,k(s) is the RMSE for method m, task q, split k, and saturation s; N_q,k(s) is the corresponding number of held-out values; M_q is the set of displayed methods eligible for task q; and K = 50.

We refer to this construction as an oracle because, although $B_1$ or $B_0$ status can be determined from source-score availability at the time of prediction, the method assigned to each task and saturation was selected after performance had been measured against the observed held-out target scores. The same 50 splits were therefore used both to identify the lowest-RMSE method for each task and to calculate the routed summary. This reuse favors methods that performed best on those particular splits and makes the reported routed RMSE an optimistic estimate of performance. The routed curve should therefore be interpreted as a best-case summary of the evaluated methods, rather than an unbiased estimate of how a prespecified or trained routing strategy would perform on new data. We therefore performed no statistical test and report no confidence interval for the combined curve.

$$q \in \{B_1, B_0\}: \quad \overline{R}_{m,q(s)} = \frac{\sum_{k=1}^{K} R_{m,q,k(s)}}{K}, \quad m_q^*(s) = arg\ min_{m \in M_q} \overline{R}_{m,q(s)}, \quad K = 50$$

$$\overline{N}_{q(s)} = \frac{\sum_{k=1}^{K} N_{q,k(s)}}{K}, \qquad RMSE_{route}(s) = \sqrt{\frac{\overline{N}_{B_1(s)}\ \overline{R}_{B_1(s)}^2 + \overline{N}_{B_0(s)}\ \overline{R}_{B_0(s)}^2}{\overline{N}_{B_1(s)} + \overline{N}_{B_0(s)}}}$$

#### Variance decomposition and reported quantities

| Symbol | Meaning |
|---|---|
| $(i, j)$ | One held-out target entry (one scored matrix cell). |
| $M_{ij}$ | Observed target score used for scoring. |
| $\widehat{M}_{ij}$ | Imputed score for the same held-out target entry. |
| $\mu_{ij}$ | Unknown noise-free target score. |
| $\varepsilon_{ij};\ \sigma_{ij}^2$ | Target measurement error and its variance; $\sigma^2$ is taken as the square of the reported target standard error (SE). |
| $r_{ij} = \widehat{M}_{ij} - M_{ij}$ | Scored residual: imputed score minus observed target score. |
| $b_{ij} = E[\widehat{M}_{ij}] - \mu_{ij}$ | Bias of the imputed score for the fixed entry, training design, and mask. |
| $\tau_{ij}^2 = Var(\widehat{M}_{ij})$ | Variance of the imputed score across hypothetical repetitions with the entry, training design, and mask held fixed. |
| $N$ | Number of scored held-out predictions. |
| $SSE;\ Q$ | Sums of $r^2$ and $\sigma^2$, respectively. |
| $MSE;\ RMSE;\ \overline{\sigma^2}$ | Mean squared residual, its square root, and mean reported measurement variance, respectively. |

| Symbol | Meaning |
|---|---|
| $T_{raw}$ | Observed MSE remaining after subtracting the mean reported measurement variance. |
| $\rho_{raw}$; $VR_{raw}$ | The remainder divided by MSE and by mean measurement variance, respectively. |
| $s_{RMSE}^2$; $RMSE\ SD = s_{RMSE}$ | Sample variance and standard deviation (SD) of the 50 aggregate split-specific RMSE values; variation across masks, not a standard error, $\tau^2$, or same-entry prediction variance. |

We did not estimate bias and prediction variance separately. In the theoretical identities below, E and Var refer to hypothetical repetitions of measurement and model fitting for a fixed held-out entry, training design, and train/test mask. Thus $\tau^2$ does not include variation from changing the mask.

Under those fixed-mask repetitions, if target measurement error is centered and the imputation is uncorrelated with target measurement error, T_raw estimates squared bias plus prediction variance; it estimates pure prediction variance only if the imputer is unbiased for every scored entry. Correlated source and target measurement errors can violate this for $B_1$; no correction was calculated.

In E[MSE], the overbars denote averages across the N scored held-out prediction events; one event is one originally observed target entry held out in one split. Table S4 combines N, the sum of squared errors (SSE), and Q over those events from the relevant maps or ordered source-to-target pairs and all 50 splits before calculating the displayed quantities.

The 50 actual Monte Carlo splits primarily vary the train/test mask. RMSE SD is the sample SD of the 50 aggregate RMSE values calculated across all scored predictions within each split; it is not a standard error or $\tau^2$, and it is not same-entry prediction variance.

$$M_{ij} = \mu_{ij} + \varepsilon_{ij}, \quad r_{ij} = \widehat{M}_{ij} - M_{ij}$$

$$E[r_{ij}^2] = b_{ij}^2 + \tau_{ij}^2 + \sigma_{ij}^2, \quad E[MSE] = \overline{b^2} + \overline{\tau^2} + \overline{\sigma^2}$$

$$MSE = \frac{SSE}{N}, \quad RMSE = \sqrt{MSE}, \quad \overline{\sigma^2} = \frac{Q}{N}$$

$$T_{raw} = MSE - \overline{\sigma^2}, \quad \rho_{raw} = \frac{T_{raw}}{MSE}, \quad VR_{raw} = \frac{T_{raw}}{\overline{\sigma^2}}$$

**Figure S1. Sensitivity of within-map PCA imputation to component number**

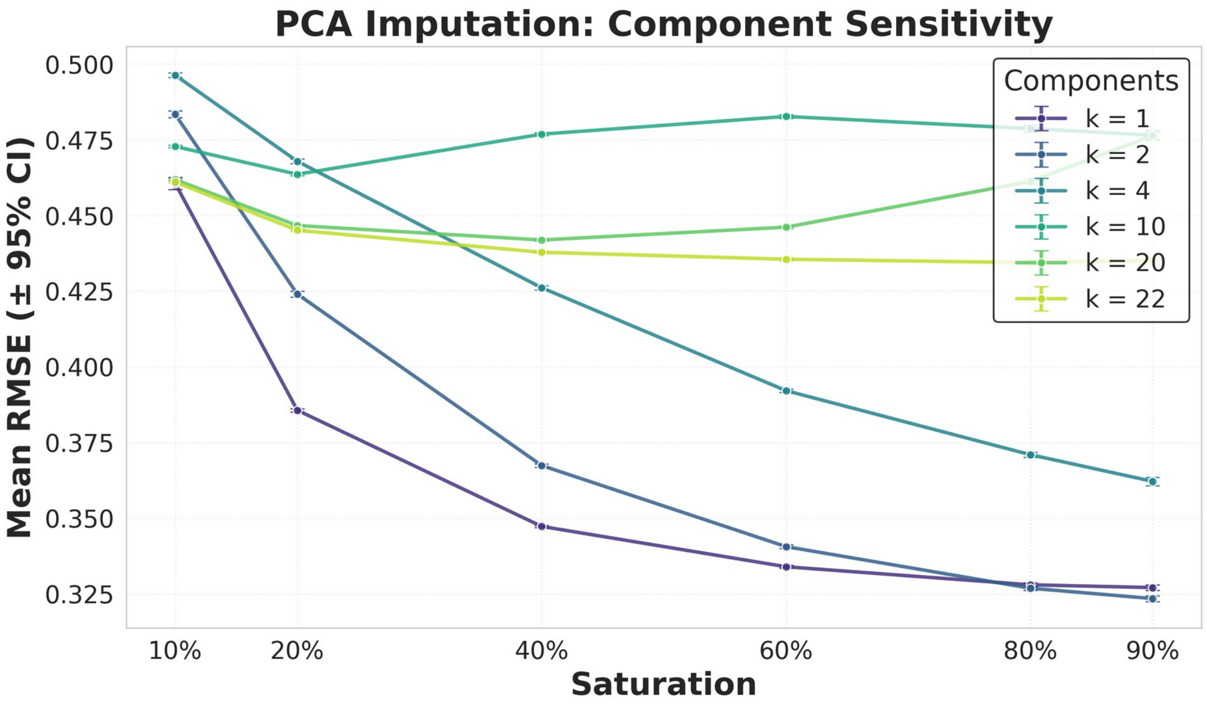


Within-map PCA reconstruction is compared for k = 1, 2, 4, 10, 20, and 22 components. Points show mean RMSE on held-out values across 50 random Monte Carlo splits, and error bars show 95% confidence intervals for the mean across splits. k = 2 is slightly lower at 90% and 80% saturation, whereas k = 1 is lower from 60% through 10% saturation; adding many components does not provide a uniformly better imputer.

### Supplemental tables

**Table S1. Source-informed RMSE at extreme target sparsity with no additional source-map masking**

#### Source-informed ($B_1$)

| Model | 90% | 60% | 20% | 1% | 0.1% |
|---|---|---|---|---|---|
| Basic Linear | 0.2811 [0.2802, 0.2820] | 0.2809 [0.2806, 0.2812] | 0.2810 [0.2809, 0.2812] | 0.2842 [0.2839, 0.2844] | 0.3212 [0.3167, 0.3256] |
| Column Mean | 0.3563 [0.3552, 0.3574] | 0.3627 [0.3623, 0.3631] | 0.4061 [0.4057, 0.4065] | 0.4542 [0.4536, 0.4548] | 0.4658 [0.4636, 0.4681] |
| DualAE | 0.2817 [0.2808, 0.2826] | 0.2939 [0.2935, 0.2942] | 0.3613 [0.3607, 0.3619] | 0.5924 [0.5920, 0.5928] | 0.6137 [0.6136, 0.6139] |
| Linear + Domain | 0.2694 [0.2685, 0.2703] | 0.2694 [0.2691, 0.2697] | 0.2699 [0.2697, 0.2700] | 0.2843 [0.2836, 0.2851] | — |
| Mixed (rand. int.) | 0.2649 [0.2640, 0.2658] | 0.2651 [0.2648, 0.2654] | 0.2661 [0.2660, 0.2662] | 0.2847 [0.2839, 0.2854] | — |
| MICE-PMM | 0.3699 [0.3691, 0.3706] | 0.3708 [0.3705, 0.3711] | 0.3723 [0.3720, 0.3727] | 0.4253 [0.4230, 0.4276] | 0.5692 [0.5620, 0.5765] |
| MICE-RF | 0.2492 [0.2483, 0.2500] | 0.2506 [0.2502, 0.2509] | 0.2556 [0.2554, 0.2558] | 0.2890 [0.2883, 0.2897] | 0.3891 [0.3860, 0.3922] |
| Mixed (rand. slope) | 0.2690 [0.2681, 0.2699] | 0.2692 [0.2689, 0.2695] | 0.2708 [0.2706, 0.2709] | 0.2878 [0.2869, 0.2886] | — |
| 1-Param Nonlinear | 0.2947 [0.2937, 0.2957] | 0.2946 [0.2942, 0.2949] | 0.2947 [0.2945, 0.2948] | 0.2969 [0.2967, 0.2971] | 0.3189 [0.3163, 0.3215] |
| SingleAE (cross-map) | 0.2773 [0.2764, 0.2782] | 0.2908 [0.2904, 0.2912] | 0.3618 [0.3612, 0.3623] | 0.5939 [0.5935, 0.5943] | 0.6140 [0.6138, 0.6141] |

*Note.* All originally observed source-map values were retained; naturally absent source values remained absent. RMSE was calculated only on source-informed ($B_1$) held-out target values.

*Note.* An em dash indicates that results were available for fewer than 95% of the 2,800 expected ordered-pair-by-split combinations (56 ordered pairs × 50 splits) at that saturation.

*Note.* Each entry gives the mean RMSE across 50 random Monte Carlo splits, followed in square brackets by a normal-approximation 95% confidence interval for that mean. Within each split, RMSE was calculated across all held-out values from the relevant maps or ordered source-to-target pairs, so maps or pairs with more scored values contribute more. The interval is $\bar{R} \pm 1.96 \times s\_R / \sqrt{50}$, where s_R is the sample SD of the 50 RMSE values. The individual RMSE values are not assumed to be normally distributed; the approximation is for their mean. The interval describes variation among the 50 masks for this fixed dataset.

*Note.* Column Mean predictions were calculated from retained target-map training scores in each split and evaluated on the same $B_1$ held-out values.

**Table S2. Method with the lowest mean RMSE by task and saturation**

#### Summary across 50 random Monte Carlo splits

| Task | Saturation | Lowest-mean method | Mean RMSE | Comparisons |
|---|---|---|---|---|
| W | 90% | PCA (k = 1) | 0.3271 | 4 |
| W | 80% | PCA (k = 1) | 0.3280 | 4 |

| Task | Saturation | Lowest-mean method | Mean RMSE | Comparisons |
|---|---|---|---|---|
| W | 60% | PCA (k = 1) | 0.3340 | 4 |
| W | 40% | PCA (k = 1) | 0.3473 | 4 |
| W | 20% | MICE-RF | 0.3803 | 4 |
| W | 10% | MICE-RF | 0.3893 | 4 |
| $B_1$ | 90% | MICE-RF | 0.2500 | 9 |
| $B_1$ | 80% | MICE-RF | 0.2518 | 9 |
| $B_1$ | 60% | MICE-RF | 0.2561 | 9 |
| $B_1$ | 40% | MICE-RF | 0.2627 | 9 |
| $B_1$ | 20% | Mixed (rand. int.) | 0.2711 | 9 |
| $B_1$ | 10% | Basic Linear | 0.2841 | 9 |
| $B_0$ | 90% | DualAE | 0.3856 | 4 |
| $B_0$ | 80% | DualAE | 0.3633 | 4 |
| $B_0$ | 60% | DualAE | 0.3546 | 4 |
| $B_0$ | 40% | DualAE | 0.3637 | 4 |
| $B_0$ | 20% | DualAE | 0.4044 | 4 |
| $B_0$ | 10% | Column Mean | 0.4501 | 4 |
| $B_1$ (no added source missingness) | 90% | MICE-RF | 0.2492 | 9 |
| $B_1$ (no added source missingness) | 60% | MICE-RF | 0.2506 | 9 |
| $B_1$ (no added source missingness) | 20% | MICE-RF | 0.2556 | 9 |
| $B_1$ (no added source missingness) | 1% | Basic Linear | 0.2842 | 9 |
| $B_1$ (no added source missingness) | 0.1% | 1-Param Nonlinear | 0.3189 | 6 |

*Note.* For each task and saturation, the listed method had the lowest mean RMSE among the methods and configurations included in the corresponding main comparison.

*Note.* The Comparisons column gives the number of contrasts between the listed method and the other included methods. Tests were two-sided unpaired Mann–Whitney U tests, with Bonferroni correction across all pairwise method comparisons within each task and saturation. Full test results are provided in the accompanying statistical CSV files.

**Table S3. Performance of the overall lowest-RMSE method within individual maps and ordered source-to-target pairs**

#### Comparisons within individual maps and ordered source-to-target pairs

| Task | Saturation | Overall lowest-RMSE method | Maps or ordered pairs | Completed comparisons | Lower mean RMSE, n/N | Significantly better, n/N | Significantly worse, n/N |
|---|---|---|---|---|---|---|---|
| W | 90% | PCA (k = 1) | 8 | 32 | 32/32 | 32/32 | 0/32 |
| W | 80% | PCA (k = 1) | 8 | 32 | 32/32 | 32/32 | 0/32 |
| W | 60% | PCA (k = 1) | 8 | 32 | 32/32 | 32/32 | 0/32 |
| W | 40% | PCA (k = 1) | 8 | 32 | 32/32 | 32/32 | 0/32 |
| W | 20% | MICE-RF | 8 | 32 | 31/32 | 30/32 | 1/32 |
| W | 10% | MICE-RF | 8 | 32 | 32/32 | 32/32 | 0/32 |
| $B_1$ | 90% | MICE-RF | 56 | 504 | 467/504 | 392/504 | 11/504 |
| $B_1$ | 80% | MICE-RF | 56 | 504 | 467/504 | 413/504 | 16/504 |
| $B_1$ | 60% | MICE-RF | 56 | 504 | 457/504 | 424/504 | 19/504 |
| $B_1$ | 40% | MICE-RF | 56 | 504 | 430/504 | 391/504 | 37/504 |
| $B_1$ | 20% | Mixed (rand. int.) | 56 | 504 | 453/504 | 342/504 | 8/504 |
| $B_1$ | 10% | Basic Linear | 56 | 504 | 440/504 | 280/504 | 7/504 |
| $B_0$ | 90% | DualAE | 56 | 224 | 224/224 | 145/224 | 0/224 |
| $B_0$ | 80% | DualAE | 56 | 224 | 224/224 | 213/224 | 0/224 |
| $B_0$ | 60% | DualAE | 56 | 224 | 224/224 | 224/224 | 0/224 |
| $B_0$ | 40% | DualAE | 56 | 224 | 224/224 | 224/224 | 0/224 |
| $B_0$ | 20% | DualAE | 56 | 224 | 224/224 | 217/224 | 0/224 |
| $B_0$ | 10% | Column Mean | 56 | 224 | 202/224 | 186/224 | 13/224 |
| $B_1$ (no added source missingness) | 90% | MICE-RF | 56 | 504 | 454/504 | 393/504 | 13/504 |
| $B_1$ (no added source missingness) | 60% | MICE-RF | 56 | 504 | 465/504 | 455/504 | 22/504 |

| Task | Saturation | Overall lowest-RMSE method | Maps or ordered pairs | Completed comparisons | Lower mean RMSE, n/N | Significantly better, n/N | Significantly worse, n/N |
|---|---|---|---|---|---|---|---|
| $B_1$ (no added source missingness) | 20% | MICE-RF | 56 | 504 | 449/504 | 446/504 | 51/504 |
| $B_1$ (no added source missingness) | 1% | Basic Linear | 56 | 504 | 409/504 | 335/504 | 45/504 |
| $B_1$ (no added source missingness) | 0.1% | 1-Param Nonlinear | 56 | 336 | 315/336 | 269/336 | 2/336 |

*Note.* For each task and saturation, the method with the lowest mean RMSE across all held-out values was compared with every other evaluated method separately within each target map for W or within each ordered source-to-target pair for $B_1$ and $B_0$.

*Note.* For example, with ten displayed $B_1$ methods (nine source-informed methods plus Column Mean) and 56 ordered pairs, nine winner-versus-competitor comparisons were made within each ordered pair, giving 56 × 9 = 504 displayed comparisons. Both methods in a test always came from the same map or ordered pair; maps and ordered pairs were not compared with one another.

*Note.* Each n/N entry reports the count out of the completed comparisons shown in that row. Lower RMSE indicates better performance.

*Note.* Significance was determined using two-sided unpaired Mann–Whitney U tests on the split-level RMSE values. P-values were Bonferroni-adjusted across all completed method-versus-method tests from all maps or ordered pairs at the same task and saturation.

*Note.* Only comparisons with results from at least 48 of 50 splits for both methods were included.

##### Table S4. Variance-decomposition results across all target-map saturation levels

*Note.* The complete table is provided as the separate Excel workbook Table_S4_variance_decomposition.xlsx. It covers all six target-map saturation levels for five W methods (Column Mean, kNN-BLOSUM, MICE-RF, PCA (k = 1), and SingleAE (within-map)); five $B_1$ methods (Basic Linear, MICE-PMM, MICE-RF, SingleAE (cross-map), and DualAE); and four $B_0$ methods (MICE-PMM, MICE-RF, SingleAE (cross-map), and DualAE).

*Note.* Results are shown separately for within-map imputation (W), source-informed between-map imputation ($B_1$), and missing-source between-map imputation ($B_0$). Saturation is the percentage of originally observed target-map values retained after imposed masking. For each method, task, and saturation, the displayed quantities were calculated after combining all scored held-out predictions from the relevant target maps or ordered source-to-target pairs and all 50 splits. Maps or ordered pairs with more scored predictions therefore contribute more. N is the total number of scored held-out predictions across the 50 splits, not the number of unique matrix entries; the same entry can be held out in more than one split. The $B_0$ results combine naturally absent source values and source values made unavailable by masking.

*Note.* RMSE = $\sqrt{(SSE / N)}$, and mean $\sigma^2$ is the mean reported target SE squared over the same scored predictions. Let $R_k$ be the aggregate RMSE from split k and $\bar{R}$ be the mean of those values across K = 50 splits. The across-split sample variance is $s_{RMSE}^2 = \Sigma(R_k - \bar{R})^2 / (K - 1)$, and its square root is $s_{RMSE} = \sqrt{s_{RMSE}^2}$. The displayed RMSE is calculated from the pooled SSE and N across all 50 splits and therefore need not equal $\bar{R}$. The variance and SD describe variation among the random masks and separately fitted models; they are not a standard error or confidence interval and not $\tau^2$ or same-entry prediction variance.

*Note.* $T_{raw}$ = MSE − mean $\sigma^2$, $VR_{raw} = T_{raw}$ / mean $\sigma^2$, and $\rho_{raw} = T_{raw}$ / MSE. Under centered and calibrated target measurement error that is uncorrelated with the prediction, $T_{raw}$ estimates squared bias plus prediction variance. It estimates prediction variance alone only if the imputer is also unbiased for every scored entry. These quantities are descriptive diagnostics. For $B_1$, correlated source and target measurement errors may violate the required covariance assumption; no correction for this possibility was applied.

**Table S5. RMSE at MTHFR Trp165 for nine source-informed methods and the target-only Column Mean reference**

#### Low-to-high folinate (wt12 → wt200)

| Method | 90% | 80% | 60% | 40% | 20% | 10% |
|---|---|---|---|---|---|---|
| SingleAE (cross-map) | 0.295716 | 0.267477 | 0.265082 | 0.316186 | 0.311811 | 0.305546 |
| DualAE | 0.290137 | 0.256179 | 0.263103 | 0.301913 | 0.315985 | 0.324666 |
| MICE-PMM | 0.486331 | 0.511178 | 0.478043 | 0.410315 | 0.505804 | 0.412005 |
| MICE-RF | 0.352991 | 0.368357 | 0.380168 | 0.402592 | 0.416905 | 0.444698 |
| Basic Linear | 0.439828 | 0.447694 | 0.450818 | 0.454074 | 0.445544 | 0.455211 |
| 1-Param Nonlinear | 0.434659 | 0.442171 | 0.445012 | 0.447988 | 0.440333 | 0.449548 |
| Linear + Domain | 0.440698 | 0.448547 | 0.451554 | 0.454797 | 0.446037 | 0.456509 |
| Mixed (rand. int.) | 0.428394 | 0.443067 | 0.445439 | 0.448692 | 0.444042 | 0.455922 |
| Mixed (rand. slope) | 0.427767 | 0.444746 | 0.445335 | 0.447230 | 0.447433 | 0.464766 |
| Column Mean | 0.373831 | 0.298740 | 0.292951 | 0.326760 | 0.343454 | 0.388966 |

#### High-to-low folinate (wt200 → wt12)

| Method | 90% | 80% | 60% | 40% | 20% | 10% |
|---|---|---|---|---|---|---|
| SingleAE (cross-map) | 0.181820 | 0.203443 | 0.230191 | 0.301001 | 0.536605 | 0.694142 |
| DualAE | 0.183359 | 0.199836 | 0.218099 | 0.234614 | 0.279884 | 0.399181 |
| MICE-PMM | 0.404611 | 0.425877 | 0.420145 | 0.379783 | 0.434412 | 0.477129 |
| MICE-RF | 0.401777 | 0.414560 | 0.431639 | 0.443861 | 0.439819 | 0.480907 |
| Basic Linear | 0.468165 | 0.465656 | 0.455123 | 0.464183 | 0.436813 | 0.474268 |
| 1-Param Nonlinear | 0.533080 | 0.530973 | 0.520103 | 0.530306 | 0.502936 | 0.533549 |
| Linear + Domain | 0.409889 | 0.406890 | 0.398588 | 0.405825 | 0.377424 | 0.426644 |
| Mixed (rand. int.) | 0.411302 | 0.407698 | 0.399853 | 0.406227 | 0.377903 | 0.426968 |
| Mixed (rand. slope) | 0.420710 | 0.417843 | 0.407399 | 0.413190 | 0.390211 | 0.442769 |
| Column Mean | 0.211622 | 0.222903 | 0.225547 | 0.221875 | 0.253874 | 0.247753 |

*Note.* Percentages are target-map saturation. For each source-informed method ($B_1$), direction, and saturation, RMSE combines all held-out non-wild-type Trp165 outcomes across 50 random splits. Column Mean was calculated from retained target-map training scores at Trp165, with the target-map-wide mean used when no Trp165 training score remained, and was evaluated on the same held-out outcomes. It is an unranked reference and was excluded from ranking the nine source-informed methods. Lower RMSE is better. No statistical tests were performed.

**Table S6. Mathematical definitions of the five source-to-target regression models**

#### Source-to-target regression models

| Model | Model formula (R syntax) | Mathematical form | Fitted parameters |
|---|---|---|---|
| Basic Linear | target ~ source | $y_i = \beta_0 + \beta_1 x_i + \varepsilon_i$ | $\beta_0$: intercept; $\beta_1$: source-to-target slope; $\sigma^2$: residual variance. |
| 1-Param Nonlinear | target ~ 1 - B*(1 - source) | $y_i = 1 - B(1 - x_i) + \varepsilon_i$ | B: only fitted mean parameter; slope = B and intercept = 1 − B, so the fitted line passes through (1, 1); $\sigma^2$: residual variance. |
| Linear + Domain | target ~ source * domain | $y_i = \alpha_{d_i} + \beta_{d_i} x_i + \varepsilon_i$ | α_d, β_d: domain-specific intercept and source-to-target slope, equivalent to the source-by-domain interaction in the R formula; $\sigma^2$: residual variance. |
| Mixed (rand. int.) | target ~ source * domain + (1 \| str_aa_mut) | $y_i = \alpha_{d_i} + \beta_{d_i} x_i + b_{0,a_i} + \varepsilon_i$ | α_d, β_d: domain-specific fixed intercept and slope; $b_0$,a: mutant-amino-acid random intercept; Var($b_0$,a): random-intercept variance; $\sigma^2$: residual variance. |
| Mixed (rand. slope) | target ~ source + domain + (1 + source \| str_aa_mut) | $y_i = \alpha_{d_i} + \beta x_i + b_{0,a_i} + b_{1,a_i} x_i + \varepsilon_i$ | α_d: domain-specific fixed intercept; β: common fixed source-to-target slope; $b_0$,a and $b_1$,a: correlated mutant-amino-acid random intercept and slope with covariance matrix Σ_b; $\sigma^2$: residual variance. |

*Note.* For entry i, x_i is the observed source-map score and y_i is the target-map score; d_i is its protein domain and a_i is the mutant-amino-acid category. The residual ε_i has mean zero and variance $\sigma^2$.

*Note.* Each model was fitted separately for each ordered source-to-target pair, split, and target-map saturation level using rows where both source and target scores were observed. Basic Linear and Linear + Domain used ordinary least squares; 1-Param Nonlinear used nonlinear least squares initialized at B = 1; the two mixed models used restricted maximum likelihood (REML).

*Note.* The mixed models use zero-mean Gaussian random effects. For prediction of a mutant-amino-acid category absent from the training data, R was called with allow.new.levels = TRUE, so its random effects were set to zero and the prediction used the fixed-effects component.
